# SOX2 terminates trophectoderm competence in inner cell mass by closing trophectoderm enhancers

**DOI:** 10.64898/2026.08.22.746397

**Authors:** Naoki Hirono, Masanori Uchikawa, Arisa Tanigawa, Takeru Fujii, Yoshiki Miyasaka, Ryo Maeda, Makoto Tachibana, Kazuki Nakao, Akihito Harada, Hiroshi Sasaki

## Abstract

During embryonic development, cellular competence to respond to differentiation signals changes dynamically. Although the mechanisms underlying competence acquisition have been extensively studied, those underlying competence loss remain unclear. In preimplantation mouse embryos, Hippo signaling shifts from regulating trophectoderm (TE) fate specification to promoting epiblast maturation. During the blastocyst stage, inner cell mass (ICM) cells lose TE competence in response to the Hippo signaling effector TEAD– YAP. Here, we show that the pioneer factor SOX2 terminates TE competence in the ICM. SOX2 binding to the TEAD–YAP-dependent TE enhancer (TEE) of the TE regulator *Gata3* induces chromatin closure, suppressing TEE responsiveness to TEAD–YAP activity. This function of SOX2 requires its interaction with the corepressor TLE4 and histone deacetylase. Similar SOX2-dependent chromatin closure also occurs around other TE genes, including the TE enhancer of another TE regulator, *Cdx2*. Thus, SOX2 terminates TE competence in ICM cells by closing Hippo signaling-responsive enhancers.

**Highlights:**

- *Sox2* is required for the loss of trophectoderm (TE) competence in the inner cell mass
- SOX2 closes the chromatin at the TEAD–YAP-dependent TE enhancer (TEE) of *Gata3*
- SOX2 exerts its repressive effects on the TEE via TLE4 and HDACs
- SOX2 terminates TE competence by closing enhancers of TE regulators and other genes

## INTRODUCTION

Embryonic development is a process in which a single zygote divides and gradually differentiates into hundreds of distinct cell types in a spatially and temporally controlled manner. Importantly, only a handful of signaling pathways are repeatedly used to regulate cell differentiation, proliferation, migration, cell death, and morphogenesis throughout development. Signal-receiving cells and tissues display context-dependent differences in competence, defined as the ability to correctly interpret signaling cues and generate appropriate cellular responses.^1^ Therefore, the timely acquisition and loss of appropriate competence for these signals are crucial for proper embryonic development.

At the molecular level, one mechanism underlying changes in competence to signaling arises from alterations in the responsiveness of target gene cis-regulatory elements (CREs), or enhancers, to downstream transcription factors in the signaling pathway.^2^ Individual genes are regulated by multiple enhancers, and distinct enhancers are selectively activated depending on the developmental stage and cell type.^3–6^ This process is governed by the accessibility of downstream transcription factors to their genomic targets.^7^ Extensive studies have revealed the mechanisms underlying the acquisition of responsiveness; pioneer factors open the chromatin at specific enhancers allowing transcription factors to gain access to their genomic targets.^8–10^ In contrast, although the loss of previous competence is equally important for the proper interpretation of signals, the mechanisms by which cells lose competence remain unknown.

To address this question, we focused on the competence shift that occurs during mouse preimplantation development. In preimplantation embryos, Hippo signaling plays two distinct roles depending on the developmental stage.^11^ Hippo signaling controls the nuclear localization of the coactivator proteins Yes-associated protein 1 (YAP1) and WW domain-containing transcription regulator 1 (WWTR1; also known as TAZ).^12,13^ Hereafter, these proteins are collectively referred to as YAP. Nuclear YAP interacts with TEA domain transcription factors (TEADs) to regulate target genes.^14–16^ At the morula stage, TEAD–YAP is activated in the outer cells and promotes trophectoderm (TE) differentiation by inducing the TE-specific transcription factor genes *Cdx2* and *Gata3.*^17,18^ TE differentiation promotes blastocyst formation, producing embryos composed of the TE and the inner cell mass (ICM). During the blastocyst stage, ICM cells progressively differentiate into pluripotent epiblast (EPI) and primitive endoderm (PrE).^19^ Once EPI fate is specified, TEAD–YAP is gradually activated. At this stage, TEAD–YAP induces pluripotency genes such as *Nanog* rather than

TE genes, thereby contributing to EPI maturation.^20,21^ Thus, the transcriptional output of TEAD–YAP shifts from TE genes to pluripotency genes. In other words, the responsiveness of target genes to TEAD–YAP changes in ICM cells. Precocious activation of TEAD–YAP in the ICM during the early blastocyst stage induces ectopic *Cdx2* expression, whereas this no longer occurs by the mid-blastocyst stage.^21,22^ Therefore, during blastocyst development, ICM cells lose the responsiveness of TE genes to TEAD–YAP, which is essential for proper TEAD–YAP-dependent EPI maturation. However, the molecular mechanisms by which TE genes lose their responsiveness to TEAD–YAP remain elusive.

In this study, we investigated the mechanisms underlying the loss of TE gene responsiveness to TEAD–YAP in ICM cells. The ICM-specific transcription factor^23,24^ and pioneer factor,^25^ SOX2, is required for the loss of TE competence in ICM cells. SOX2 forms a complex with TLE4 and histone deacetylases (HDACs), binds to the TEAD–YAP-dependent enhancer of the TE regulator gene *Gata3*, and reduces enhancer accessibility. Similar mechanisms also appear to operate at *Cdx2* and several other TE genes. Thus, binding of the SOX2–TLE4–HDAC complex closes the chromatin at TEAD–YAP-regulated enhancers of TE regulators and other TE genes, thereby causing the loss of TE competence in ICM cells.

## RESULTS

### GATA3 as a sensitive marker for TE differentiation competence of ICM cells

At the morula stage, TEAD–YAP promotes TE fate specification by inducing the TE-specific transcription factors CDX2 and GATA3.^17,18,26^ The ICM of early blastocyst stage embryos retains TE differentiation competence, as demonstrated by the regeneration of miniature blastocysts from isolated ICMs^17,27^ and by ectopic CDX2 expression following experimental activation of TEAD–YAP.^21,22^ To determine when ICM cells lose TE competence, we first examined whether experimental activation of TEAD–YAP at different developmental stages induces ectopic expression of the TE-specific genes *Cdx2* and *Gata3* in the ICM. TEAD– YAP was activated by promoting YAP nuclear localization through treatment of blastocysts with the LATS1/2 inhibitor (LATSi^28^) during three developmental intervals: 0–12, 12–24, and 24–36 h after blastocoel formation (Figure 1A). The 0-h and 24-h time points correspond to the early and mid-blastocyst stages, respectively, under *in vitro* culture conditions. Embryos treated during the 0–12 h interval showed strong CDX2 and GATA3 expression in most ICM cells (Figures 1B–1D). In embryos treated during the 12–24 h interval, only a small fraction of ICM cells expressed CDX2 and GATA3 (Figures 1B-1D). Embryos treated during the 24–36 h interval did not express either CDX2 or GATA3 in the ICM (Figures 1B–1D). These results suggest that the responsiveness of *Cdx2* and *Gata3* to TEAD–YAP is maintained in the ICM during the early blastocyst stage but is lost by the mid-blastocyst stage, consistent with previous findings for *Cdx2.*^21,22^

**Figure 1.**
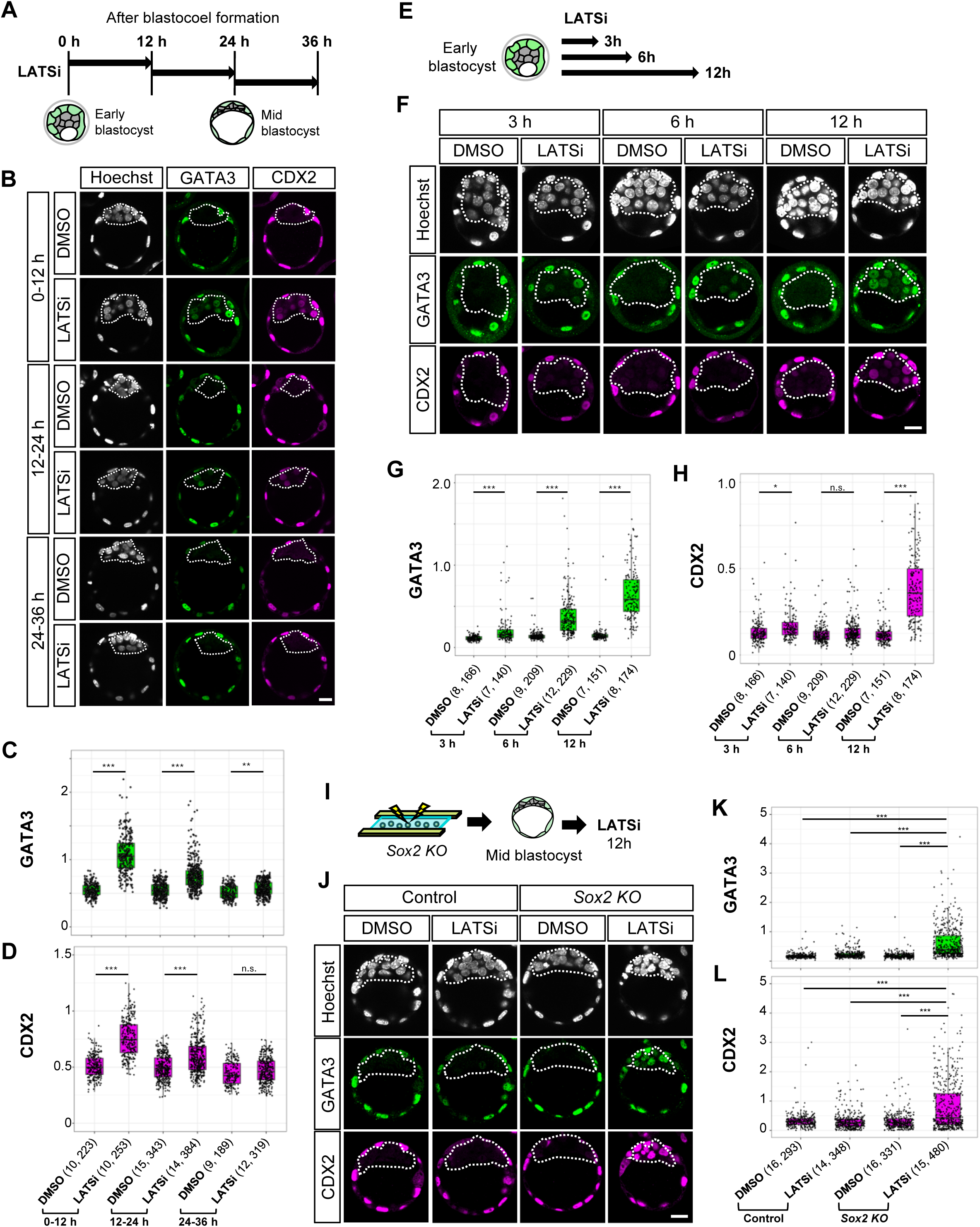
SOX2 expression is required for the loss of responsiveness of *Gata3* and *Cdx2* to TEAD–YAP in ICM cells. (A) Schematic of the experimental design. (B) Representative images of GATA3 and CDX2 expression in embryos treated with LATSi for 12 h at the indicated time points. Dashed lines indicate the ICM. Scale bar, 20 μm. (C, D) Quantification of GATA3 (C) and CDX2 (D) signal intensities in ICM cells shown in (B). Each dot represents an individual cell. The numbers following each group indicate the sample sizes (N, number of embryos; n, number of cells). \*\**p* < 0.01, \*\*\**p* < 0.001 (two-way ANOVA followed by Tukey’s multiple comparisons test); n.s., not significant. (E) Schematic of the experimental design. (F) Representative images of GATA3 and CDX2 expression in embryos treated with LATSi for the indicated time durations. Dashed lines indicate the ICM. Scale bar, 20 μm. (G, H) Quantification of GATA3 (G) and CDX2 (H) signal intensities in ICM cells shown in (F). Each dot represents an individual cell. The numbers following each group indicate the sample sizes (N, number of embryos; n, number of cells). \**p* < 0.05; \*\*\**p* < 0.001 (two-way ANOVA followed by Tukey’s multiple comparisons test); n.s., not significant. (I) Schematic of the experimental design. (J) Representative images of GATA3 and CDX2 expression in control and *Sox2 KO* embryos treated with LATSi or DMSO (control). Dashed lines indicate the ICM. Scale bar, 20 μm. (K, L) Quantification of GATA3 (K) and CDX2 (L) signal intensities in ICM cells shown in (J). Each dot represents an individual cell. The numbers following each group indicate the sample sizes (N, number of embryos; n, number of cells). \*\*\**p* < 0.001 (two-way ANOVA followed by Tukey’s multiple comparisons test).

To determine which TE regulator better reflects the TE differentiation competence of ICM cells, we compared the induction kinetics of CDX2 and GATA3 following TEAD– YAP activation. Embryos were treated with LATSi for 3, 6, or 12 h beginning at blastocoel formation (Figure 1E). GATA3 expression in ICM cells increased after 3 h of treatment and continued to increase through 12 h (Figures 1F and 1G). In contrast, CDX2 expression showed little increase during the first 6 h but was eventually upregulated in all ICM cells after 12 h of treatment (Figures 1F and 1H). Therefore, *Gata3* responds to TEAD–YAP earlier than *Cdx2* and is a more sensitive indicator of TE differentiation competence. In addition to their different induction kinetics, CDX2 and GATA3 also differ in evolutionary conservation. Although both CDX2 and GATA3 are expressed in the TE of human and bovine blastocysts, only GATA3 is expressed in morulae and early blastocysts under the control of YAP.^29,30^ CDX2 expression begins only at the mid-blastocyst stage.^31,32^ Based on its rapid induction kinetics and broader evolutionary conservation, we conclude that GATA3 is a better marker for assessing the TE differentiation competence of ICM cells in response to TEAD–YAP.

### SOX2 is required for the loss of *Gata3* responsiveness to TEAD–YAP activity

To elucidate the mechanisms by which ICM cells lose *Gata3* responsiveness to TEAD–YAP, we first searched for upstream regulators of *Gata3*. We reasoned that the acquisition of ICM identity would suppress the TE competence of ICM cells. SOX2 is the only transcription factor specifically expressed in inner cells as early as the 16-cell stage and is required to establish the pre-pluripotency state of the ICM in early blastocysts.^23,24,33^ These characteristics make SOX2 a strong candidate for an upstream regulator. To determine whether SOX2 is involved in the loss of *Gata3* responsiveness to TEAD–YAP, we generated *Sox2* knockout (KO) embryos by genome editing in zygotes, as previously described,^21,34^ and treated them with LATSi beginning at the mid-blastocyst stage, when *Gata3* responsiveness is normally lost (Figure 1I). Whereas LATS inhibition did not induce GATA3 expression in the ICM of control (*DsRed* KO) embryos, it induced ectopic GATA3 expression in the ICM of *Sox2* KO embryos (Figures 1J and 1K). These results suggest that SOX2 is required for the loss of *Gata3* responsiveness to TEAD–YAP.

### Identification of the TE enhancer of *Gata3*

Analysis of *Sox2* KO embryos led us to hypothesize that the loss of responsiveness is caused by epigenetic changes at TEAD–YAP-dependent enhancers of *Gata3* and that these changes are regulated by SOX2. To investigate this hypothesis, we first searched for enhancer candidates co-regulated by TEAD–YAP and SOX2 using published epigenomic datasets. Candidate enhancers were expected to meet the following criteria. (1) Enhancers of competent genes should be in a poised state with open chromatin. In wild-type embryos, *Gata3* responsiveness is present at the early blastocyst stage but is lost by the mid-blastocyst stage. Accordingly, ATAC-seq peaks in ICM cells are present in early blastocysts but markedly reduced in late blastocysts (wild-type ATAC-seq tracks in Figures 2Ai and S1Ai).^33^ (2) Because *Gata3* responsiveness is maintained in *Sox2* KO embryos at the mid-blastocyst stage, ATAC-seq peaks in the ICM of late-stage *Sox2* KO blastocysts should remain comparable to those in early blastocysts (*Sox2* KO ATAC-seq tracks in Figures 2Aii and S1Aii).^33^ (3) Because SOX2 is required for the change in responsiveness, SOX2 binding should be detected by CUT&RUN in ICM cells (SOX2 CUT&RUN tracks in Figures 2Aiii and S1Aiii).^33^ (4) Because *Gata3* is regulated by TEAD4–YAP in the TE, TEAD4 binding should be detectable by chromatin immunoprecipitation followed by sequencing (ChIP-seq) in trophoblast stem cells (TSCs), which resemble the TE (TEAD4 ChIP-seq track in Figures 2Aiv and S1Aiv).^35^ We identified two adjacent peaks that satisfied all four criteria. These peaks were located downstream of the *Gata3* locus, +56,048 to +61,466 bp from the transcription start site (TSS) (Figures 2A and S1A). The upstream and downstream peaks were designated P1and P2, respectively, and the combined region encompassing both peaks was termed P1+P2.

**Figure 2.**
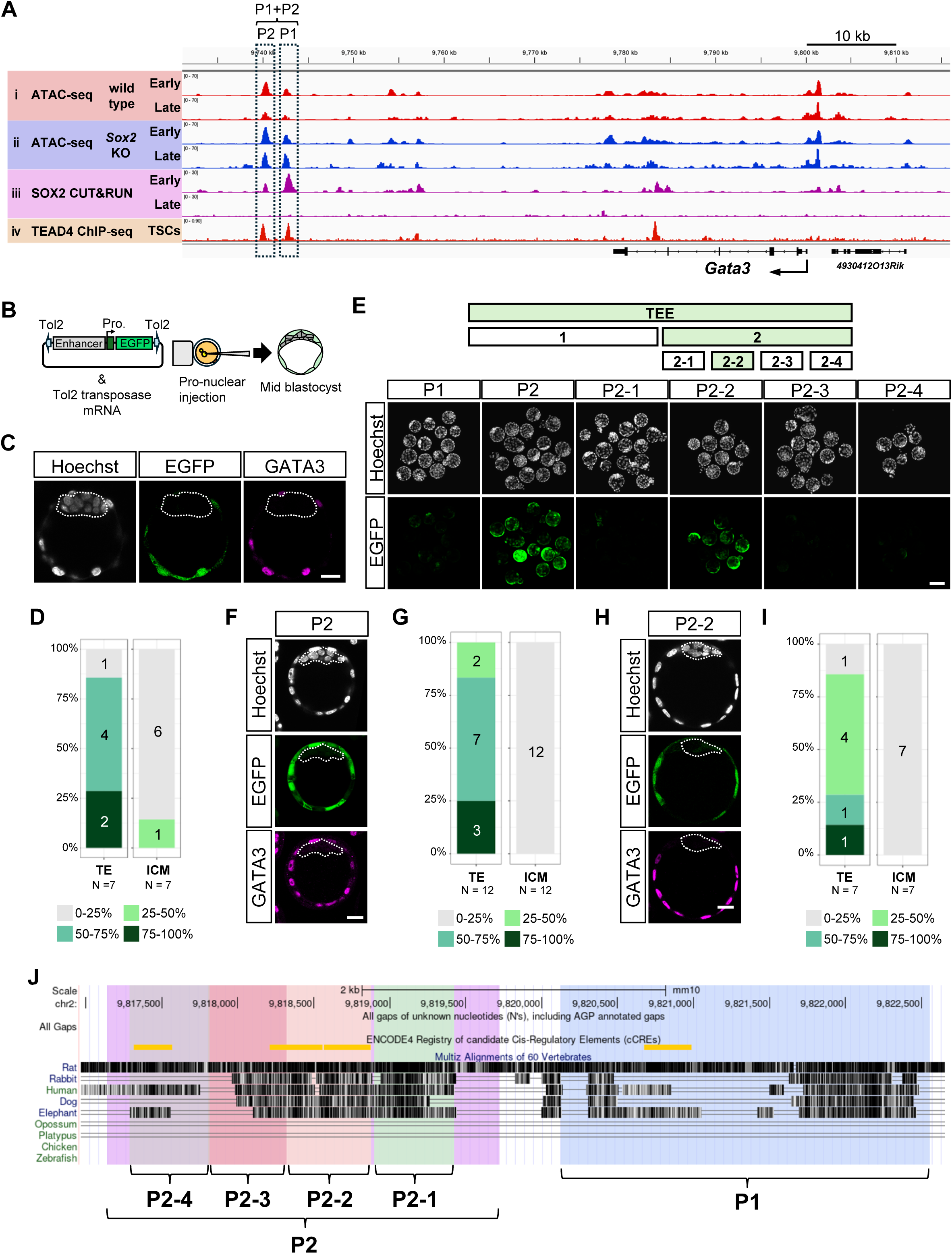
A TE-specific enhancer of *Gata3* is located downstream of the *Gata3* locus. (A) IGV snapshots of the genomic region surrounding the *Gata3* locus. Early, ICM of early (E3.5) blastocysts; Late, ICM of late (E4.5) blastocysts; TSCs, trophoblast stem cells. (B) Schematic of the transgenic reporter assay in F0 embryos. (C) Representative images of EGFP and GATA3 expression in P1+P2 transgenic embryos. Dashed lines indicate the ICM. Scale bar, 20 μm. (D) Graph summarizing EGFP expression frequencies in P1+P2 transgenic embryos. Embryos were categorized according to the proportion of EGFP-positive cells in each cell type. The numbers in each column indicate the number of embryos in each category. (E) Representative images of EGFP expression in TEE-subfragment transgenic embryos. Scale bar, 100 μm. (F) Representative images of EGFP and GATA3 expression in P2 transgenic embryos. Dashed lines indicate the ICM. Scale bar, 20 μm. (G) Graph summarizing EGFP expression frequencies in P2 transgenic embryos. (H) Representative images of EGFP fluorescence and GATA3 immunofluorescence in P2-2 transgenic embryos. Dashed lines indicate the ICM. Scale bar, 20 μm. (I) Graph summarizing EGFP expression frequencies in P2-2 transgenic embryos. (J) UCSC Genome Browser^77^ snapshots of the TEE showing the cross-species conservation of each segment. Yellow bars indicate ENCODE candidate cis-regulatory elements. See also Figure S1.

To determine whether P1+P2 functions as an enhancer, we performed F0 transgenic enhancer assays in mouse embryos. A reporter plasmid, in which P1+P2 was placed upstream of an enhanced green fluorescent protein (EGFP) reporter cassette and the entire construct was flanked by inverted terminal repeats (ITRs) of the *Tol2* transposon, was co-injected with *Tol2* transposase mRNA into the pronuclei of zygotes to generate transgenic embryos (Figure 2B). The resulting embryos were examined for EGFP expression at the mid-blastocyst stage (Figure 2B). Most P1+P2-transgenic embryos showed TE-specific EGFP expression (Figures 2C and 2D). Therefore, P1+P2 possesses TE-specific enhancer activity and is hereafter referred to as the TE enhancer (TEE).

To identify the core element of TEE, we examined the activities of P1 and P2 individually. Whereas P1-transgenic embryos showed no EGFP expression, P2-transgenic embryos exhibited TE-specific EGFP expression, indicating that P2 plays the dominant role (Figures 2E–2G). To further define the core element of TEE, P2 was divided into four subfragments (P2-1, P2-2, P2-3, and P2-4). Only P2-2 (551 bp) showed TE-specific EGFP expression, whereas the other fragments were inactive (Figures 2E, 2H, and 2I), suggesting that P2-2 constitutes the core element of TEE.

Consistent with these findings, P2-2 contains two ENCODE candidate cis-regulatory elements (cCREs), characterized by high DNase hypersensitivity and H3K27ac enrichment (Figure 2J, yellow bars).^36^ The P2-2 region also exhibits high evolutionary conservation across placental mammals (Figure 2J). In contrast, orthologous sequences corresponding to P2-2 or TEE were not detected in non-placental mammals (opossum and platypus) or lower vertebrates (chicken and zebrafish) (Figure 2J). These results suggest that TEE is associated with implantation and/or placental development and that P2-2 represents the evolutionarily conserved core of the enhancer.

### TEE recapitulates *Gata3* responsiveness to TEAD–YAP in the ICM

Because TEE recapitulates the TE-specific expression of *Gata3*, we next examined whether TEE also recapitulates the responsiveness of endogenous *Gata3* to TEAD–YAP in the ICM. For this purpose, TEE-transgenic embryos were treated with LATSi for 24 h, starting at either the early or mid-blastocyst stage, to activate TEAD–YAP and assess EGFP expression in the ICM. Embryos treated from the early blastocyst stage showed EGFP and GATA3 expression in the ICM (Figures 3A and 3B). In contrast, embryos treated from the mid-blastocyst stage showed neither EGFP nor GATA3 expression in the ICM (Figures 3C and 3D). A similar stage-dependent responsiveness to TEAD–YAP was observed in P2-2-transgenic embryos (Figures 3E–3H). Thus, TEE and its core element, P2-2, recapitulate the stage-dependent responsiveness of *Gata3* to TEAD–YAP in the ICM. These results suggest that TEE functions as the enhancer that mediates TEAD–YAP-dependent activation of *Gata3*, with P2-2 serving as its core element.

**Figure 3.**
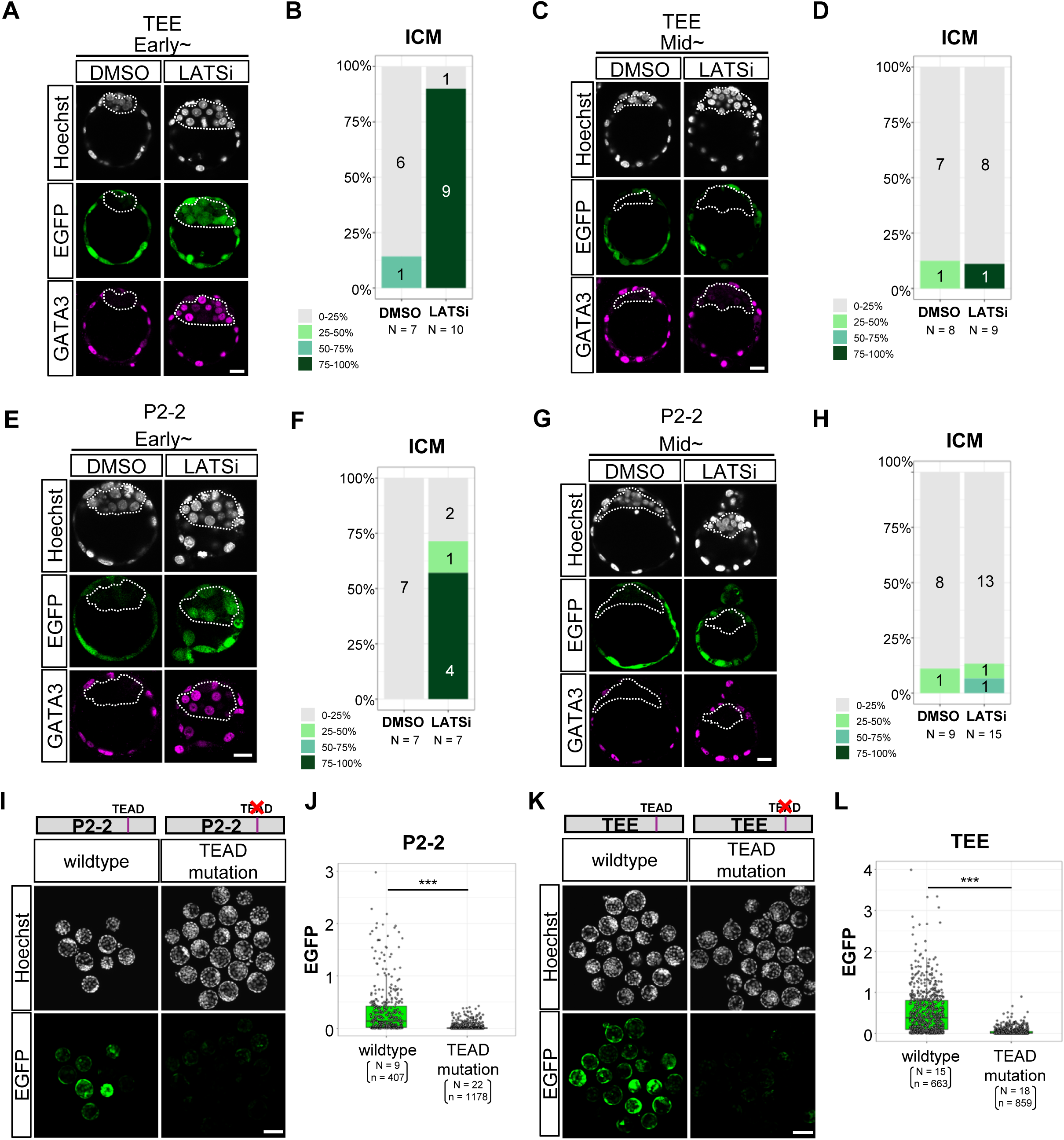
The TEE recapitulates the responsiveness of endogenous *Gata3* to TEAD–YAP in ICM cells. (A) Representative images of EGFP and GATA3 expression in TEE transgenic embryos treated with LATSi beginning at the early blastocyst stage. Dashed lines indicate the ICM. Scale bar, 20 μm. (B) Graph summarizing EGFP expression frequencies in the ICM of the transgenic embryos shown in (A). Embryos were categorized according to the proportion of EGFP-positive cells. The numbers in each column indicate the number of embryos in each category. (C) Representative images of EGFP and GATA3 expression in TEE transgenic embryos treated with LATSi beginning at the mid-blastocyst stage. Dashed lines indicate the ICM. Scale bar, 20 μm. (D) Graph summarizing EGFP expression frequencies in the ICM of the transgenic embryos shown in (C). (E) Representative images of EGFP and GATA3 expression in P2-2 transgenic embryos treated with LATSi beginning at the early blastocyst stage. Dashed lines indicate the ICM. Scale bar, 20 μm. (F) Graph summarizing EGFP expression frequencies in the ICM of the transgenic embryos shown in (E). (G) Representative images of EGFP and GATA3 expression in P2-2 transgenic embryos treated with LATSi beginning at the mid-blastocyst stage. Dashed lines indicate the ICM. Scale bar, 20 μm. (H) Graph summarizing EGFP expression frequencies in the ICM of the transgenic embryos shown in (G). (I) Representative images of EGFP expression in wild-type and TEAD motif-mutated P2-2 transgenic embryos at the mid-blastocyst stage. Scale bar, 100 μm. (J) Box plots showing EGFP signal intensities in TE cells of the embryos shown in (I). Each dot represents an individual cell. The number of embryos analyzed (N) is indicated above each column. \*\*\**p* < 0.001 (two-tailed unpaired Student’s *t*-test). (K) Representative images of EGFP fluorescence in wild-type and TEAD motif-mutated TEE transgenic embryos. Scale bar, 100 μm. (L) Box plots showing EGFP signal intensities in TE cells of the embryos shown in (K). Each dot represents an individual cell. \*\*\**p* < 0.001 (two-tailed unpaired Student’s *t*-test). See also Figure S2.

Because TEE/P2-2 is activated by TEAD–YAP in the ICM (Figures 3A and 3B) and is bound by TEAD4 in TSCs (Figure 2Aiv), we hypothesized that TEE/P2-2 is directly regulated by TEAD–YAP. To test this hypothesis, we mutated the consensus TEAD-binding motif (5′-CATTCC-3′)^37^ within P2-2 (Figures S2A and S2B) and examined its enhancer activity in embryos. Transgenic embryos carrying the TEAD motif-mutated P2-2 construct showed no EGFP expression in the TE at the mid-blastocyst stage (Figures 3I and 3J). Mutation of the same TEAD motif within the entire TEE likewise abolished EGFP expression (Figures 3K and 3L), indicating that TEAD–YAP binding to P2-2 is essential for TEE activation. These results suggest that TEE is directly regulated by TEAD–YAP and faithfully recapitulates *Gata3* responsiveness to TEAD–YAP. Given that chromatin accessibility at the TEE decreases in the ICM between the early and late blastocyst stages (Figure 2Ai), this chromatin closure may underlie the loss of *Gata3* responsiveness to TEAD–YAP during blastocyst development.

### SOX2 binding to TEE promotes the loss of *Gata3* responsiveness to TEAD–YAP in the ICM

Because *Sox2* is required for the loss of *Gata3* responsiveness to TEAD–YAP (Figures 1J and 1K) and SOX2 binds to TEE at the early blastocyst stage (Figures 2Aiii and S1Aiii),^33^ we reasoned that SOX2 binding to TEE promotes the loss of *Gata3* responsiveness to TEAD– YAP. To test this hypothesis, we mutated the consensus SOX2-binding motif (5′-AACAATT-3′) ^38,39^ within P2-2 (Figures S3A and S3B) and examined its enhancer activity in embryos. Transgenic embryos carrying the SOX2 motif-mutated P2-2 construct were treated with LATSi beginning at the mid-blastocyst stage, when endogenous *Gata3* no longer responds to TEAD–YAP. In untreated embryos, EGFP expression in the TE was unaffected by the SOX2 motif mutation (Figure 4A). Upon LATSi treatment, however, EGFP was induced in the ICM of embryos carrying the SOX2 motif-mutated P2-2 construct, but not in embryos carrying the wild-type P2-2 construct (Figures 4A and 4B). The absence of GATA3 expression in the ICM confirmed that endogenous *Gata3* had lost responsiveness (Figure 4A). These results suggest that SOX2 binding to P2-2 is required for the loss of P2-2 responsiveness to TEAD–YAP in the ICM.

**Figure 4.**
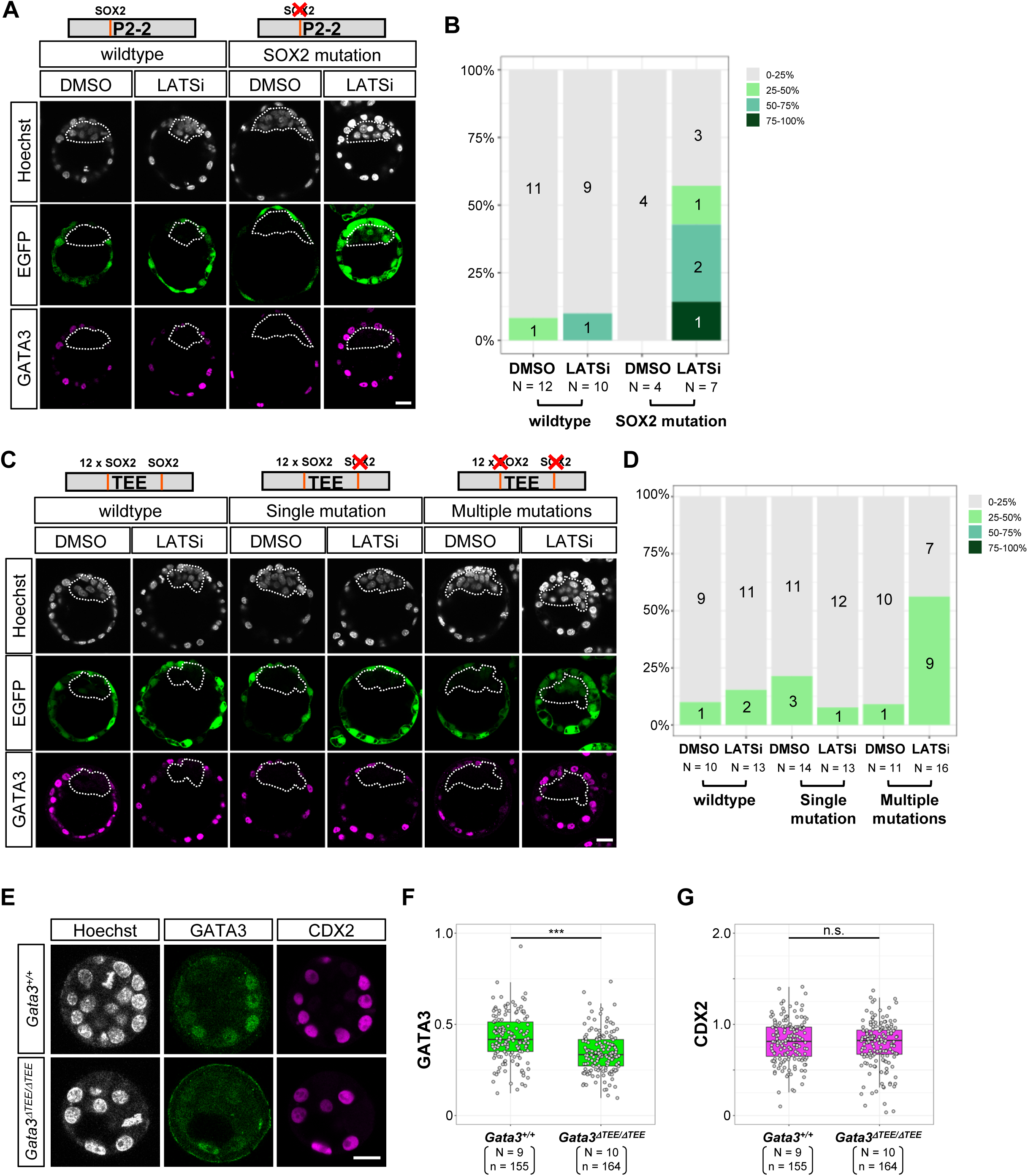
SOX2 bound at multiple sites within the TEE cooperatively mediates the loss of responsiveness to TEAD–YAP in the ICM. (A) Representative images of EGFP and GATA3 expression in wild-type and SOX2 motif-mutated P2-2 transgenic embryos treated with LATSi beginning at the mid-blastocyst stage. Dashed lines indicate the ICM. Scale bar, 20 μm. (B) Graph summarizing EGFP expression frequencies in the ICM of the transgenic embryos shown in (A). Embryos were categorized according to the proportion of EGFP-positive cells. The numbers in each column indicate the number of embryos in each category. (C) Representative images of EGFP and GATA3 expression in wild-type and SOX2 motif-mutated TEE transgenic embryos treated with LATSi. Dashed lines indicate the ICM. Scale bar, 20 μm. (D) Graph summarizing EGFP expression frequencies in the ICM of the transgenic embryos shown in (C). (E) Representative images of GATA3 and CDX2 expression in wild-type (*Gata3^+/+^*), and *Gata3^ΔTEE/ΔTEE^* embryos at the early blastocyst stage. (F, G) Box plots showing GATA3 (F) and CDX2 (G) signal intensities in TE cells of the embryos shown in (E). Each dot represents an individual cell. \*\*\**p* < 0.001 (two-way ANOVA followed by Tukey’s multiple comparisons test); n.s., not significant. See also Figure S3.

Given the importance of the SOX2 motif in P2-2 for the loss of its responsiveness, we next asked whether this motif is also critical for regulation of the entire TEE. The TEE carrying a mutation in this SOX2 motif did not express EGFP in the ICM upon LATSi treatment (Figures 4C and 4D, single mutation), indicating that this SOX2 motif alone is dispensable for the loss of TEE responsiveness to TEAD–YAP. SOX2 CUT&RUN analysis showed stronger SOX2 binding to P1 (Figure 2A iii),^33^ and sequence analysis of P1 revealed the presence of 12 consecutive and/or overlapping SOX2 motifs (Figure S3C). Therefore, we mutated all SOX2 motifs present in P1 and P2-2 (multiple mutations) (Figure S3D). TEE carrying mutations in multiple SOX2-motifs showed EGFP expression in the ICM upon LATSi treatment (Figures 4C and 4D, multiple mutations). These results suggest that SOX2 proteins bound at multiple sites within the TEE cooperatively promote the loss of responsiveness to TEAD–YAP in the ICM, thereby extinguishing TE enhancer activity. Given that the reduction in chromatin accessibility at the TEE in the ICM is *Sox2*-dependent (Figure 2A)^33^, multiple SOX2 proteins bound to the TEE likely cooperate to promote chromatin closure in the ICM, thereby causing the loss of *Gata3* responsiveness to TEAD– YAP.

### TEE is required for strong expression of GATA3

Given that TEE is regulated by TEAD–YAP and SOX2, we next investigated whether this enhancer is required for *Gata3* expression in the TE. For this purpose, we established a TEE deletion mouse line, *Gata3^ΔTEE^*, by genome editing in zygotes (Figure S3E). Genomic sequencing of the mutant allele confirmed deletion of the entire TEE region (Figure S3F). Early blastocyst-stage embryos obtained by intercrossing *Gata3^ΔTEE/+^* mice were analyzed. Although GATA3 expression was not completely abolished in *Gata3^ΔTEE/ΔTEE^* embryos, its expression level was significantly lower than that in *Gata3 ^+/+^* embryos (Figures 4E and 4F). CDX2 expression was unaffected by deletion of the TEE (Figures 4E and 4G), indicating that the reduction was specific to *Gata3* and directly attributable to the absence of TEE. These results suggest that *Gata3* expression in the TE is regulated by multiple enhancers, among which TEE plays a major role.

### SOX2 suppresses *Gata3* responsiveness to TEAD–YAP by forming a complex with TLE4 and HDACs

Although SOX2 is widely recognized as a pioneer factor that binds DNA and promotes chromatin opening,^25,40^ our results suggest that SOX2 binding instead promotes enhancer closure. We therefore next investigated the mechanism by which SOX2 closes chromatin. Because SOX2 functions as both a transcriptional activator and a repressor in neural stem cells, and its repressor activity is mediated through interactions with the TLE family corepressors TLE3 and TLE5,^41^ we hypothesized that a similar repressive mechanism operates during chromatin closure in the ICM. To test this hypothesis, we first analyzed the expression of *Tle* family genes during blastocyst development. Analysis of published scRNA-seq data revealed that *Tle4* is the predominant *Tle* gene expressed in the ICM at the early blastocyst stage and that its expression is maintained in the EPI of late blastocysts (Figure 5A).^42^

**Figure 5.**
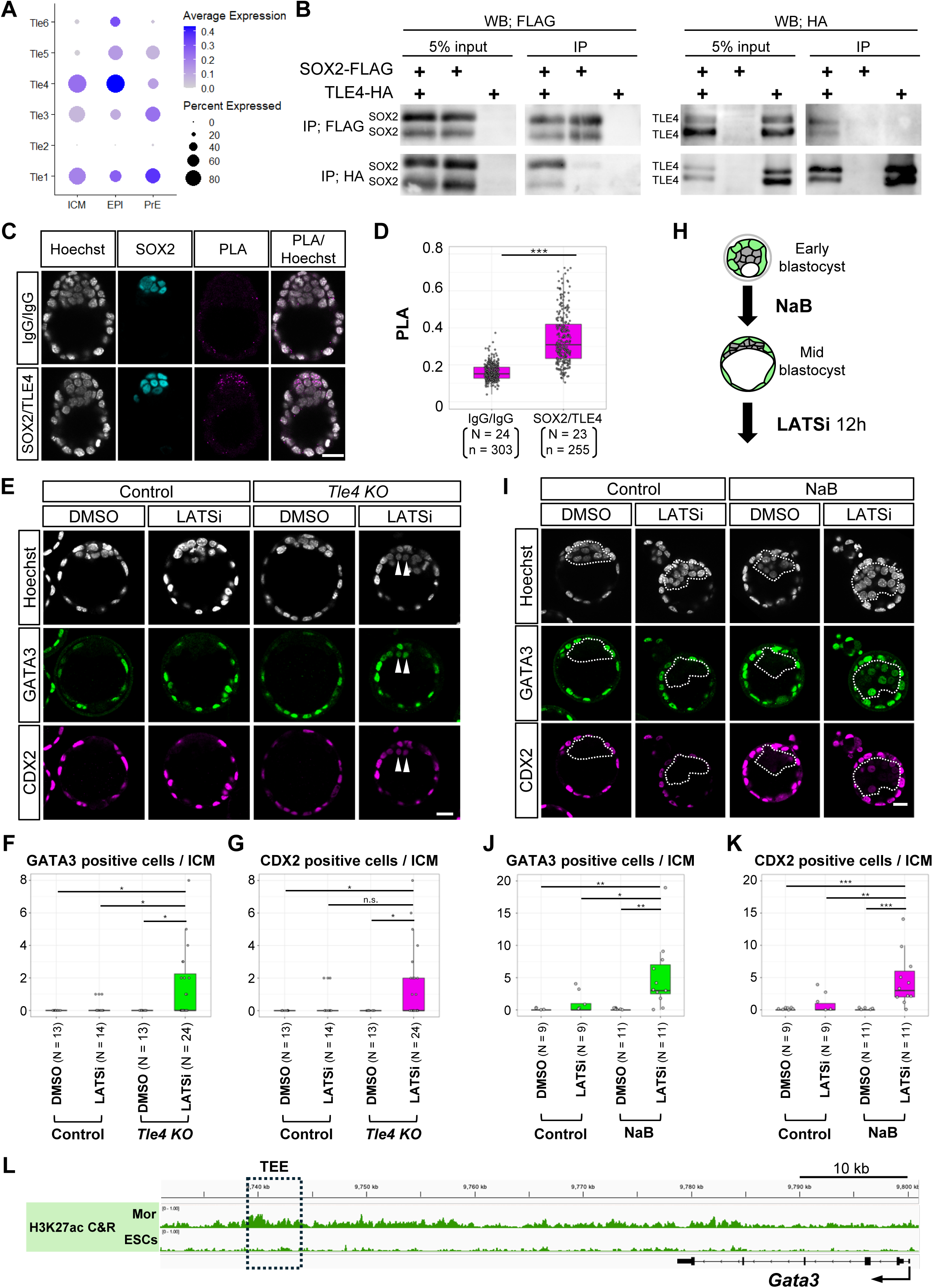
SOX2-dependent loss of GATA3 responsiveness in ICM cells is mediated by TLE4 and HDACs. (A) Dot plot showing the expression of *Tle* family genes in blastocysts^42^. ICM, ICM of early (E3.5) blastocysts. (B) Co-immunoprecipitation of SOX2-FLAG and TLE4-HA in HEK293T cells. (C) Representative images of PLA detecting SOX2 and TLE4 interactions in E4.0 embryos. Scale bar, 20 μm. (D) Box plots showing PLA signal intensities in SOX2-positive ICM cells of the embryos shown in (C). Each dot represents an individual cell. \*\*\**p* < 0.001 (two-tailed unpaired Student’s *t*-test). (E) Representative images of GATA3 and CDX2 expression in control and *Tle4* KO embryos treated with LATSi beginning at the mid-blastocyst stage. Arrowheads indicate GATA3- and CDX2-positive cells. Scale bar, 20 μm. (F, G) Box plots showing the numbers of GATA3-positive (F) and CDX2-positive (G) cells in the ICM of the embryos shown in (E). Each dot represents an individual embryo. \**p* < 0.05 (two-way ANOVA followed by Tukey’s multiple comparisons test); n.s., not significant. (H) Schematic of the experimental design. (I) Representative images of GATA3 and CDX2 expression in control and NaB-treated embryos. Dashed lines indicate the ICM. Scale bar, 20 μm. (J, K) Box plots showing the numbers of GATA3-positive (J) and CDX2-positive (K) cells in the ICM of the embryos shown in (I). Each dot represents an individual embryo. \**p* < 0.05; \*\**p* < 0.01; \*\*\**p* < 0.001 (two-way ANOVA followed by Tukey’s multiple comparisons test). (L) IGV snapshots of the genomic region surrounding the *Gata3* locus showing H3K27ac CUT&RUN signals in morulae and ESCs. The dashed box indicates the TEE. Mor, morula. See also Figure S4.

To determine whether SOX2 interacts with TLE4, we first performed co-immunoprecipitation assays using overexpressed proteins and found that FLAG-tagged SOX2 (SOX2-FLAG) co-precipitated with HA-tagged TLE4 (TLE4-HA) in HEK293T cells, and vice versa (Figure 5B). These results suggest that SOX2 and TLE4 form a complex in cultured cells. Next, we investigated whether this interaction also occurs *in vivo* using a proximity ligation assay (PLA) for SOX2 and TLE4 at the mid-blastocyst stage. Strong PLA signals were observed in SOX2-positive ICM cells, suggesting an interaction between SOX2 and TLE4 *in vivo* (Figures 5C and 5D). Finally, to determine whether TLE4 is involved in the loss of *Gata3* responsiveness to TEAD–YAP, we generated *Tle4* KO embryos by CRISPR/Cas9-mediated genome editing in zygotes. Immunofluorescence confirmed the absence of TLE4 protein in all embryos at the mid-blastocyst stage (N = 10/10), indicating 100% knockout efficiency (Figure S4A). Loss of *Tle4* did not significantly affect preimplantation development, as indicated by the normal total cell number at the late blastocyst stage (Figures S4A and S4B). Whereas activation of TEAD–YAP by LATSi beginning at the mid-blastocyst stage did not induce GATA3 expression in the ICM of control embryos, the same treatment induced GATA3 expression in a subset of ICM cells in *Tle4* KO embryos (Figures 5E and 5F). These results suggest that TLE proteins are required for the SOX2-mediated loss of *Gata3* responsiveness to TEAD–YAP in the ICM, with TLE4 playing a major role.

TLE proteins interact with histone deacetylases (HDACs) through the N-terminal GP domain of TLE proteins.^43,44^ This raised the possibility that the SOX2–TLE4 complex promotes chromatin closure by deacetylating histones through HDACs. To test this hypothesis, embryos were first treated with sodium butyrate (NaB), an inhibitor of most class I and class IIa HDACs,^45^ from the early to the mid-blastocyst stage and were then treated with LATSi beginning at the mid-blastocyst stage to activate TEAD–YAP (Figure 5H). In contrast to untreated control embryos, NaB-treated embryos expressed GATA3 in the ICM following LATSi treatment (Figures 5I and 5J). These results suggest that HDAC activity is required for the loss of *Gata3* responsiveness to TEAD–YAP. Supporting this conclusion, comparison of H3K27ac CUT&RUN profiles between morulae and embryonic stem cells (ESCs), which resemble the early ICM and EPI, respectively, revealed that H3K27ac peaks in the TEE region were reduced in ESCs (Figures 5L and S4C).^46^ Taken together, these results are consistent with a model in which the SOX2–TLE4–HDAC complex bound to the TEE reduces chromatin accessibility by promoting H3K27 deacetylation, thereby contributing to the loss of *Gata3* responsiveness to TEAD–YAP.

### Chromatin remodeling by SOX2 binding terminates the responsiveness of some TE genes to TEAD–YAP

Analysis of the *Gata3* TEE revealed that SOX2 restricts chromatin accessibility at the TE enhancer to terminate responsiveness to TEAD–YAP. We next investigated whether this mechanism also operates at other TE genes. To address this question, we isolated ICMs from control and *Sox2* KO embryos treated with LATSi beginning at the mid-blastocyst stage for 12 h and compared their gene expression profiles by RNA-seq (Figure 6A). A total of 284 and 88 genes were significantly upregulated in *Sox2* KO and control ICMs, respectively (Figures 6B and 6D, Tables S1 and S2). Gene ontology (GO) enrichment analysis of differentially expressed genes (DEGs) revealed enrichment of TE-associated categories (embryonic placenta development, establishment or maintenance of cell polarity, and actin filament organization) among genes upregulated in the *Sox2* KO ICM (Figure 6C; Table S3). The upregulated genes included TE marker genes (*Gata3, Cdx2, Krt8, Krt18,* and *Id2*)^18,23,47–49^ and apical domain genes (*Cldn4* and *Shroom3*)^50,51^ (Figure 6B). Indeed, 46% (130/284) of the genes upregulated in the *Sox2* KO ICM were specifically expressed in E3.5 and/or E4.5 TE,^52^ whereas only 12.5% (11/88) of those upregulated in the wild-type ICM were TE genes (Figures 6D and 6E; Tables S1 and S2). Notably, among TE-specific transcription factors, *Gata3* and *Cdx2*, which function downstream of TEAD4–YAP,^17,18^ were induced, whereas *Eomes* and *Elf5*, which function in later trophoblast maturation,^48,53^ were not induced (Figure 6B). These results support the hypothesis that SOX2 restricts the responsiveness of TEAD– YAP target genes in the TE.

**Figure 6.**
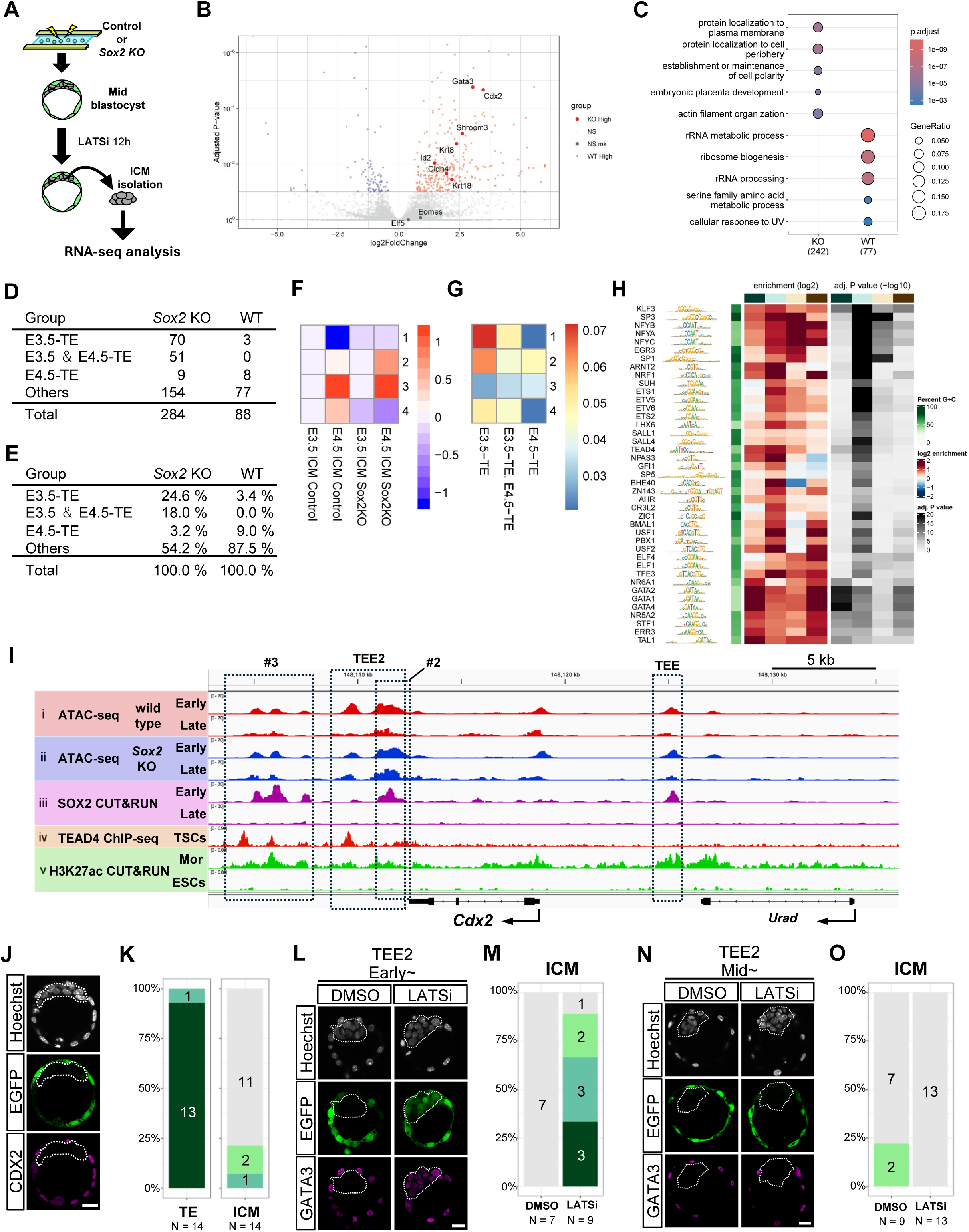
SOX2 restricts the responsiveness of multiple TE genes to TEAD–YAP by remodeling chromatin accessibility. (A) Schematic of the experimental design. Individual isolated ICMs were subjected to RNA-seq. (B) Volcano plot of RNA-seq data from control and *Sox2 KO* ICMs treated with LATSi. Red and blue dots indicate genes that were significantly upregulated and downregulated, respectively, in *Sox2 KO* ICMs. Adjusted *p* value (*p*adj) = 0.1. (C) Gene ontology enrichment analysis of DEGs between wild-type and *Sox2 KO* ICMs treated with LATSi. (D, E) Tables summarizing the numbers (D) and percentages (E) of DEGs that overlap with genes specifically expressed in the E3.5 and/or E4.5 TE.^52^ (F) Clustering analysis of ATAC-seq peaks. (G) Proportions of TE marker genes^52^ among genes nearest to high ATAC-seq peaks that overlap with high SOX2 CUT&RUN peaks in E3.5 control ICMs, shown for each ATAC-seq peak cluster. The values indicate the Jaccard coefficients. (H) Transcription factor motif enrichment analysis of high ATAC-seq peaks that overlap with high SOX2 CUT&RUN peaks in E3.5 control ICMs, shown for each ATAC-seq peak cluster. The columns represent clusters 1, 2, 3, and 4 from left to right. (I) IGV snapshots of the genomic region surrounding the *Cdx2* locus. Early, ICMs of E3.5 blastocysts; Late, ICMs of E4.5 blastocysts; Mor, morulae. Dashed boxes indicate enhancers. (J) Representative images of EGFP and CDX2 expression in *Cdx2* TEE2 transgenic embryos. Dashed lines indicate the ICM. Scale bar, 20 μm. (K) Graph summarizing EGFP expression frequencies in *Cdx2* TEE2 transgenic embryos. Embryos were categorized according to the proportion of EGFP-positive cells in each cell type. The numbers in each column indicate the number of embryos in each category. (L) Representative images of EGFP and CDX2 expression in *Cdx2* TEE2 transgenic embryos treated with LATSi beginning at the early blastocyst stage. Dashed lines indicate the ICM. Scale bar, 20 μm. (M) Graph summarizing EGFP expression frequencies in the embryos shown in (L). (N) Representative images of EGFP and CDX2 expression in *Cdx2* TEE2 transgenic embryos treated with LATSi beginning at the mid-blastocyst stage. Dashed lines indicate the ICM. Scale bar, 20 μm. (O) Graph summarizing EGFP expression frequencies in the embryos shown in (N). See also Figure S5.

To further test this hypothesis, we examined whether genomic regions surrounding genes upregulated in the *Sox2* KO ICM exhibit SOX2-dependent changes in chromatin accessibility and SOX2/TEAD4 binding similar to those observed at the *Gata3* TEE (Figure 2A). Notably, the downstream region of *Cdx2* fulfilled these criteria, and a 3-kb fragment containing two adjacent ATAC-seq peaks (TEE2) exhibited TE enhancer activity in transgenic embryos (Figures 6I–6K). TEE2 is distinct from the two previously identified TE enhancers located upstream of the TSS (TEE)^26^ and further downstream (#3),^54^ indicating that *Cdx2* expression in the TE is regulated by multiple enhancers. Furthermore, TEE2 responded to TEAD–YAP in the ICM at the early blastocyst stage but not at the mid-blastocyst stage (Figures 6L–6O). These results suggest that genomic regions meeting the criteria described above possess TE enhancer activity and that *Cdx2* responsiveness to TEAD–YAP is also regulated by SOX2. Consistent with this conclusion, ectopic CDX2 expression in the ICM was also observed in *Sox2* KO, *Tle4* KO, and NaB-treated embryos following LATSi treatment beginning at the mid-blastocyst stage (Figures 1J, 1L, 5E, 5G, 5I, and 5K), further supporting the hypothesis that the SOX2–TLE4–HDAC complex restricts chromatin accessibility at the *Cdx2* enhancer.

To examine the generality of SOX2-dependent regulation of chromatin accessibility at TE genes, ATAC-seq peaks were clustered according to developmental stage and *Sox2* genotype. Cluster 1 exhibited a *Sox2*-dependent reduction in chromatin accessibility at E4.5 (late blastocysts) (Figure 6F). To characterize these candidate regions, highly accessible ATAC-seq peaks overlapping strong SOX2 CUT&RUN peaks were selected for further analysis. E3.5 TE genes were the most enriched genes associated with Cluster 1 peaks (Figure 6G). Notably, TEAD4-binding motifs were enriched in this cluster (Figure 6H). These genomic regions are therefore likely to function as TE enhancers whose responsiveness to TEAD–YAP in the ICM is restricted by the SOX2–TLE4–HDAC complex. Consistent with this idea, Cluster 1 peaks overlapping with TEAD4 ChIP-seq peaks in TSCs were identified near or within TE-specific genes, including *Id2*, *Krt8/18*, *Cldn4*, and *Shroom3* (Figure S5A). Taken together, restriction of TE enhancer responsiveness to TEAD–YAP by the SOX2–TLE4–HDAC complex in key TE regulators (*Gata3* and *Cdx2*) as well as several TE-specific genes is likely to underlie the loss of TE competence in ICM cells.

## DISCUSSION

During development, timely changes in cellular developmental competence are essential for proper cell differentiation and embryonic development. During the transition from the morula to EPI, TEAD–YAP target genes change in coordination with cell differentiation. TE commitment of outer cells occurs between the late 16-cell and early 32-cell stages, whereas ICM cells retain TE differentiation competence until the 64-cell stage.^22^ After the 64-cell stage, EPI fate is specified in the ICM.^19^ TEAD–YAP is then activated in EPI-specified cells and induces *Nanog* expression to promote EPI maturation.^21^ Thus, the absence of TE competence in EPI-specified cells is essential for proper EPI development regulated by TEAD–YAP. Here, we show that SOX2 terminates TE competence in ICM cells. Our findings support a model in which the SOX2–TLE4–HDAC complex binds to the accessible TEE of the TE regulator *Gata3*,^18,55^ removes H3K27ac, and closes the chromatin. Consequently, TEE becomes inaccessible to TEAD–YAP, and *Gata3* loses responsiveness (Figure 7). Similar SOX2-dependent chromatin closure also appears to occur at TEAD–YAP-regulated enhancers of another TE regulator, *Cdx2*^47,48^ and of several genes involved in epithelial integrity and cytoskeletal organization (*Krt8/18*,^18,56,57^ *Cldn4*,^58^ and *Shroom3*^51^, etc.). Therefore, SOX2 terminates TE competence in ICM cells by suppressing the responsiveness of key transcriptional regulators of TE differentiation, as well as several genes involved in TE maturation, to TEAD–YAP (Figure 7).

**Figure 7.**
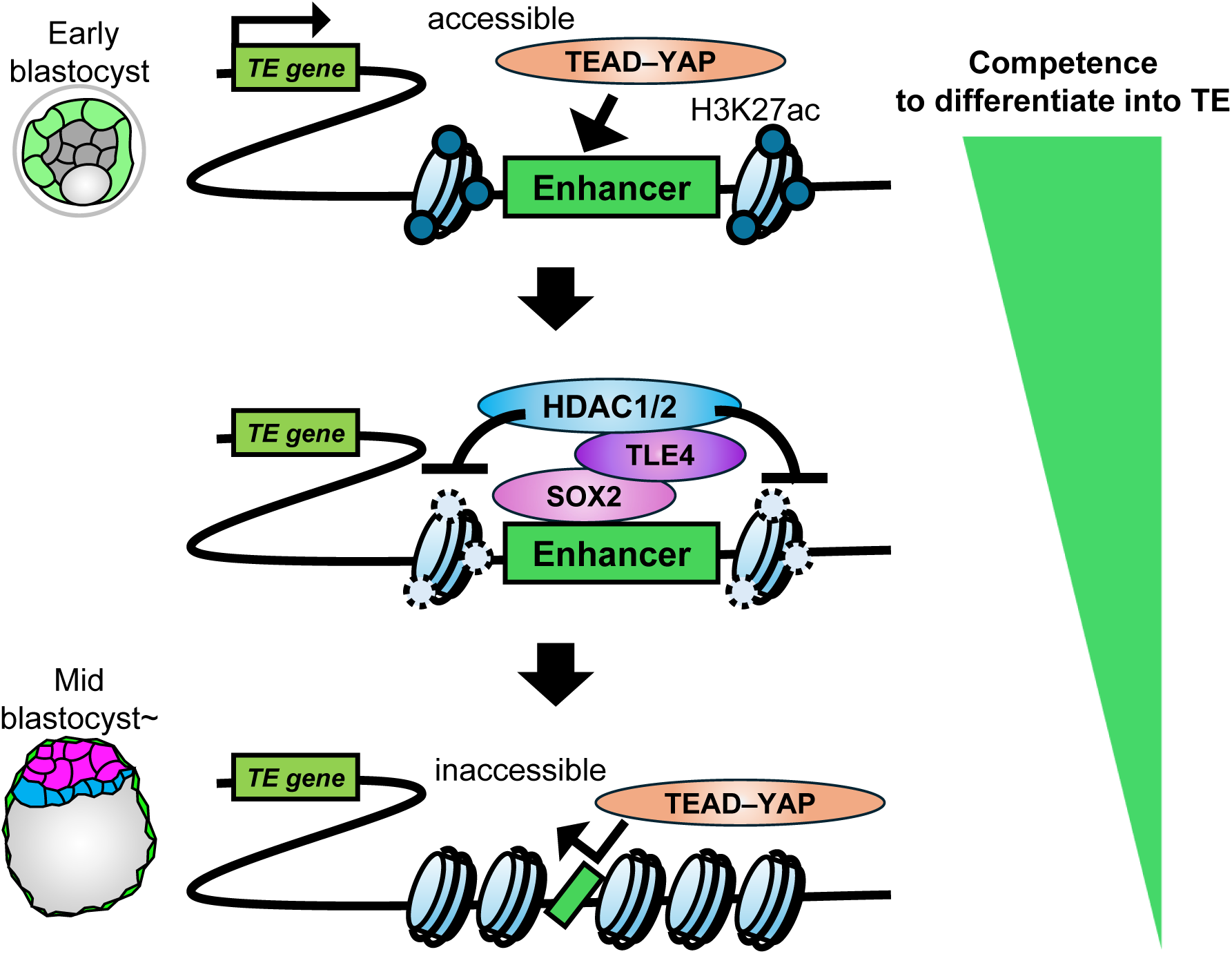
Model for the SOX2-dependent loss of TE competence during ICM development. See the Discussion for details. See also Figure S6.

SOX2 is a pioneer factor^25^ capable of initiating chromatin opening.^59^ During preimplantation development, however, SOX2 exhibits distinct modes of enhancer binding while regulating the pluripotency network. In the E4.5 EPI, SOX2 exhibits "pioneer binding," in which it opens enhancer chromatin.^33^ In contrast, in the early (E3.5) ICM, SOX2 exhibits "settler binding," in which it binds pre-accessible enhancers of EPI and PrE genes and establishes the pre-pluripotency state of the ICM, characterized by weak coexpression of EPI and PrE lineage transcription factors.^33^ Our study identifies an additional mode of SOX2 binding in the early ICM. The TE enhancers of *Gata3* and *Cdx2* are already accessible from the 8-cell stage (Figures S6A and S6B).^60^ SOX2 binds these pre-accessible TE enhancers and closes the chromatin, thereby terminating their responsiveness to TEAD–YAP. We refer to this mode of SOX2 binding as "closer binding." Because *Sox2* mutant embryos exhibit upregulation of TE gene transcripts in the ICM,^33,61^ closer binding may also contribute to suppressing TE gene expression in the early ICM. However, its primary function appears to be terminating enhancer responsiveness at the chromatin level, because ectopic expression of TE proteins (CDX2, GATA3, and EOMES) is not detected in the ICM of *Sox2* mutants^24^ (Figure 1), and these proteins become detectable only after TEAD–YAP activation in the early ICM of wild-type embryos or in the mid-ICM of *Sox2* mutant embryos (Figures 1B and 1J). Thus, both settler binding and closer binding contribute to establishing the pre-pluripotency chromatin state, thereby preparing cells for subsequent differentiation into the EPI or PrE.

The chromatin-closing activity of SOX2 is mediated through its interaction with the Groucho-related corepressor TLE4 and HDACs. Recruitment of TLE4 by SOX2 likely promotes the formation of locally closed chromatin through the direct interaction of TLE4 with chromatin, as previously shown for the TLE3–FOXA1/HES1 complex.^62^ TLE/Groucho family proteins interact with HDACs through their GP domain,^43^ and HDAC-mediated removal of H3K27ac promotes chromatin closure,^63,64^ as demonstrated for Groucho–HDAC-dependent increase in nucleosome density in *Drosophila*.^65^ Because H3K27ac is a hallmark of active enhancers,^64^ the TEE in the early ICM is likely in a primed state, and removal of H3K27ac may convert it into a poised state. Defining the precise epigenetic transition accompanying competence loss will require further analysis of chromatin modifications in the ICM. HDAC1/2-dependent repression of TSC genes (*Cdx2*, *Krt8/18*, etc.) has also been reported in ESCs,^45^ suggesting that HDACs contribute not only to the loss of enhancer responsiveness but also to maintenance of the repressed state of TE genes. More broadly, HDAC-dependent restriction of chromatin accessibility and developmental competence has been reported in other developmental contexts,^66^ suggesting that HDAC-mediated regulation of developmental competence may represent a more general mechanism.

Recent studies have shown that some pioneer factors also mediate gene silencing through diverse mechanisms.^67^ Although these repressive functions have been described primarily in artificial settings, such as cellular reprogramming and cell fate switching,^68–71^ our study demonstrates that a similar mechanism operates during normal embryonic development to terminate developmental competence. The mechanism by which SOX2 silences TE enhancers in the ICM resembles the inactivation of somatic enhancers during induced pluripotent stem cell (iPSC) reprogramming by the OSK factors (OCT4, SOX2, and KLF4), in which HDAC recruitment promotes chromatin closure.^68^ Matsui et al. also reported a repressive function of FOXA in preventing alternative cell fates during endoderm development using *in vitro* differentiation model.^72^ Although this function resembles that described here, the underlying mechanism is distinct because it depends on interactions with PRDM1/14 rather than the TLE-HDAC complex.

The TE enhancer (TEE2) identified downstream of *Cdx2* is distinct from the two previously identified enhancers, TEE and enhancer #3,^26,54^ indicating that *Cdx2* expression is regulated by multiple enhancers. Notably, TEE2 encompasses the previously tested #2 region, which lacked TE enhancer activity when examined in isolation (Figure 6I).^54^ This observation raises the possibility that the two adjacent ATAC-seq peaks function cooperatively as a single enhancer, analogous to the P1 and P2 elements of the *Gata3* TEE. Likewise, homozygous deletion of the *Gata3* TEE did not completely abolish GATA3 expression in the TE, suggesting that *Gata3* is also regulated by multiple TE enhancers. Although the additional TE enhancers controlling *Gata3* remain to be identified, redundant enhancers regulating the key TE regulators *Cdx2* and *Gata3* may confer robustness to TE development, as has been reported for many developmental genes in mice and *Drosophila*.^73–75^

The role of TEAD–YAP in TE fate specification is conserved among mammals, including humans and cattle.^29,30^ Whether the mechanism underlying the loss of TE competence identified in this study is similarly conserved, however, remains unknown. In mice, SOX2 expression is restricted to the ICM from the early blastocyst stage onward, whereas in humans and cattle, SOX2 is initially expressed in both the ICM and TE and becomes restricted to the ICM/EPI only at the late blastocyst stage.^29,30^ This temporal difference in SOX2 expression raises two possibilities. First, the loss of TE competence in humans and cattle may be mediated by other transcription factors and/or cofactors specifically expressed in the early ICM. Alternatively, TE competence may be lost only after SOX2 expression becomes restricted to the ICM. The latter possibility is consistent with the observation that human TE cells retain the ability to generate ICM cells until the full blastocyst stage (Bl3, Day 5),^76^ whereas mouse TE cells become committed by the late 32-cell (early blastocyst) stage.^22^ Thus, differences in the timing of developmental plasticity loss may underlie the distinct patterns of SOX2 expression observed among mammalian species. Comparative analyses of TE competence and SOX2 expression across mammalian embryos will help clarify the evolutionary conservation and divergence of this mechanism.

## RESOURCE AVAILABILITY

### Lead contact

Further information and requests for resources and reagents should be directed to and will be fulfilled by the lead contact Hiroshi Sasaki.

### Materials availability

The materials generated in this study are available upon request.

### Data and code availability

- RNA-seq data have been deposited in the Gene Expression Omnibus (GEO) under accession number GSE331074.
- This study does not report original code.
- Any additional information required to reanalyze the data reported in this paper is available from the lead contact upon request.

## ACKNOWLEDGMENTS

We thank Masakazu Hashimoto for advice on ICM isolation and RNA-seq; Koichi Kawakami for the Tol2 system plasmids; Hitoshi Niwa for *Sox2* cDNA; the NGS core facility at the Research Institute for Microbial Diseases of The University of Osaka for RNA sequencing and data processing. The mouse work carried out in the Sasaki lab was supported by JSPS KAKENHI (grant numbers JP19H04778, JP20H03261, JP21H05288, JP26K020167 to H.S.), and JST SPRING (grant number JPMJSP2138 to H.N.). Bioinformatics work carried out in the Harada lab was supported by JSPS KAKENHI (grant numbers JP21H05292, JP23K27087, JP23H02394, and JP26H01535 to A.H.; JP26K18286 to T.F.), AMED-CREST (grant number JP23gm1810008 to A.H.), the MEXT Promotion of Development of a Joint Usage/Research System Project: Pan-Omics DDIRC, MRCI for High Depth Omics, CURE (grant number JPMXP1323015486 for MIB), RIIT, and AMRC in Kyushu University to A.H., and research grants from the Uehara Memorial Foundation and the Shinnihon Foundation of Advanced Medical Treatment Research to A.H.

## AUTHOR CONTRIBUTIONS

N.H. and H.S. conceptualized the project. H.S. supervised the project. N.H., M.U., A.H., A.T., Y.M., M.T. and H.S. designed the experiments. N.H. performed most mouse experiments, IGV analysis of published epigenomics data, and co-IP experiments with the help of M.U. and R.M. H.S. performed reanalysis of published scRNA-seq data. A.T., Y.M., and K.N. generated the *Gata3^ΔTEE^* mouse line. M.U. analyzed *Gata3^ΔTEE/ΔTEE^* embryos. N.H. prepared samples for RNA-seq. T.F. and A.H. performed RNA-seq data analysis and reanalysis of published epigenomic datasets. N.H., M.U., A.H., and H.S. prepared the figures. The manuscript was written by N.H. and H.S., with contributions from all authors.

## STAR METHODS

### KEY RESOURCES TABLE

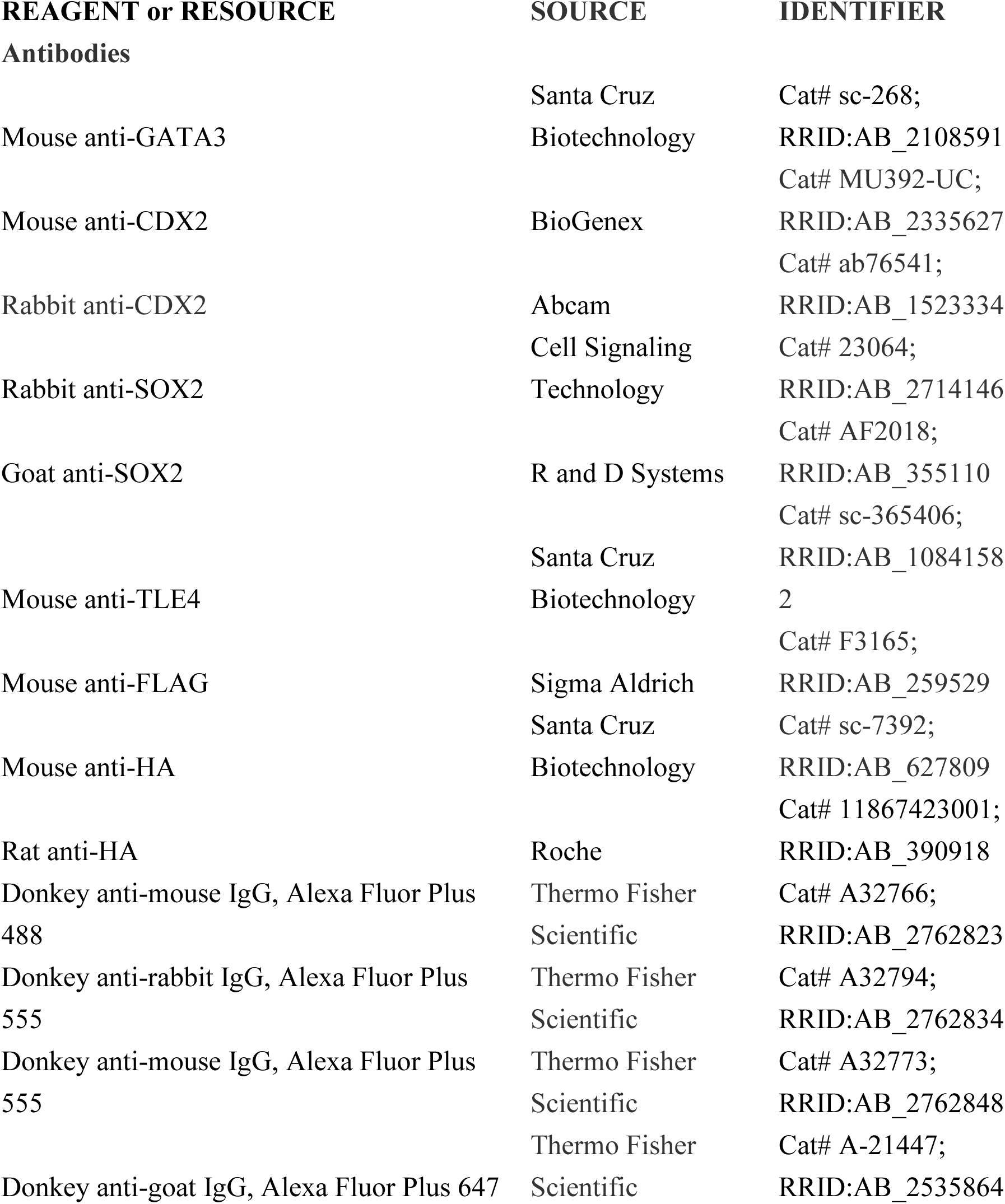
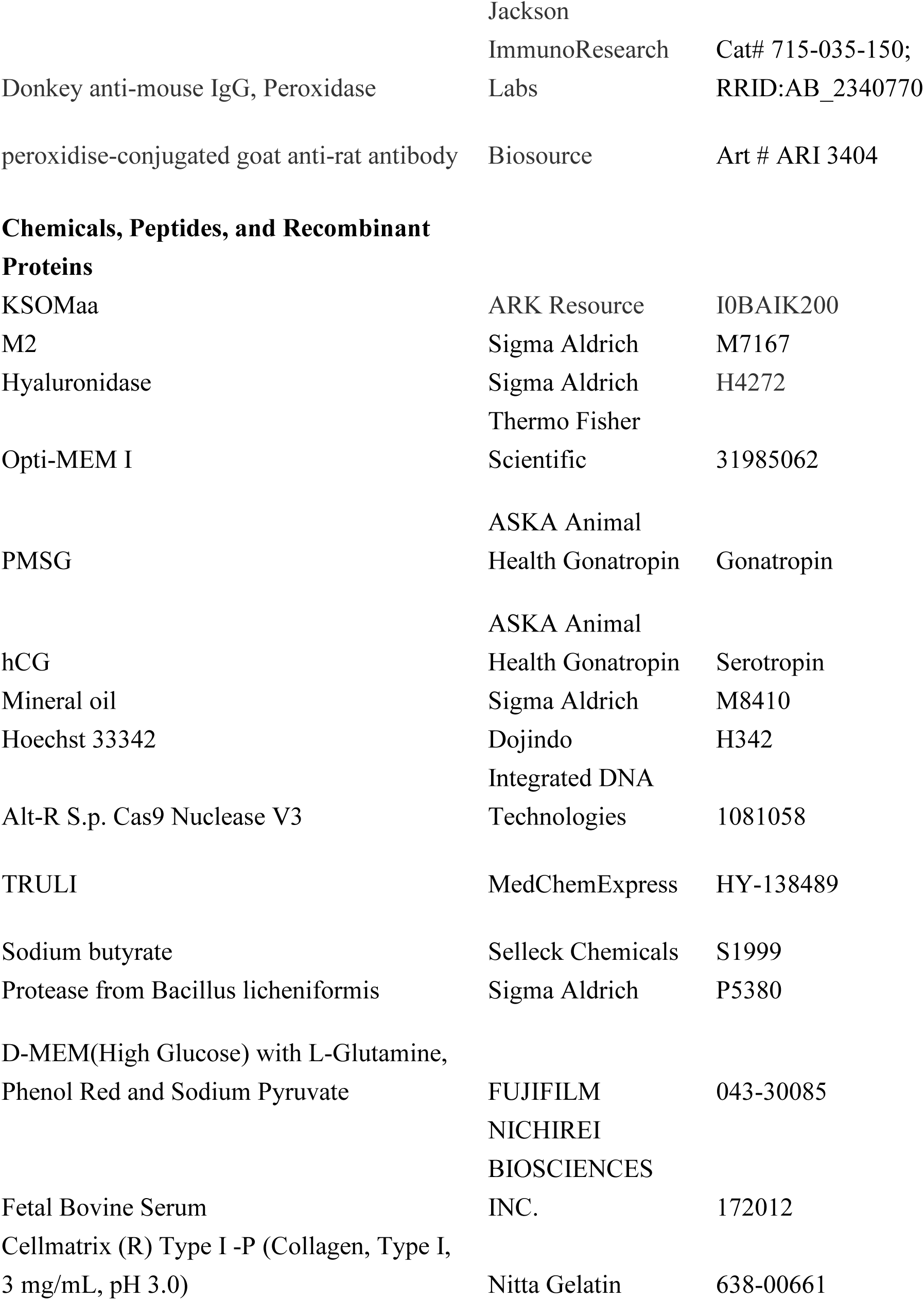
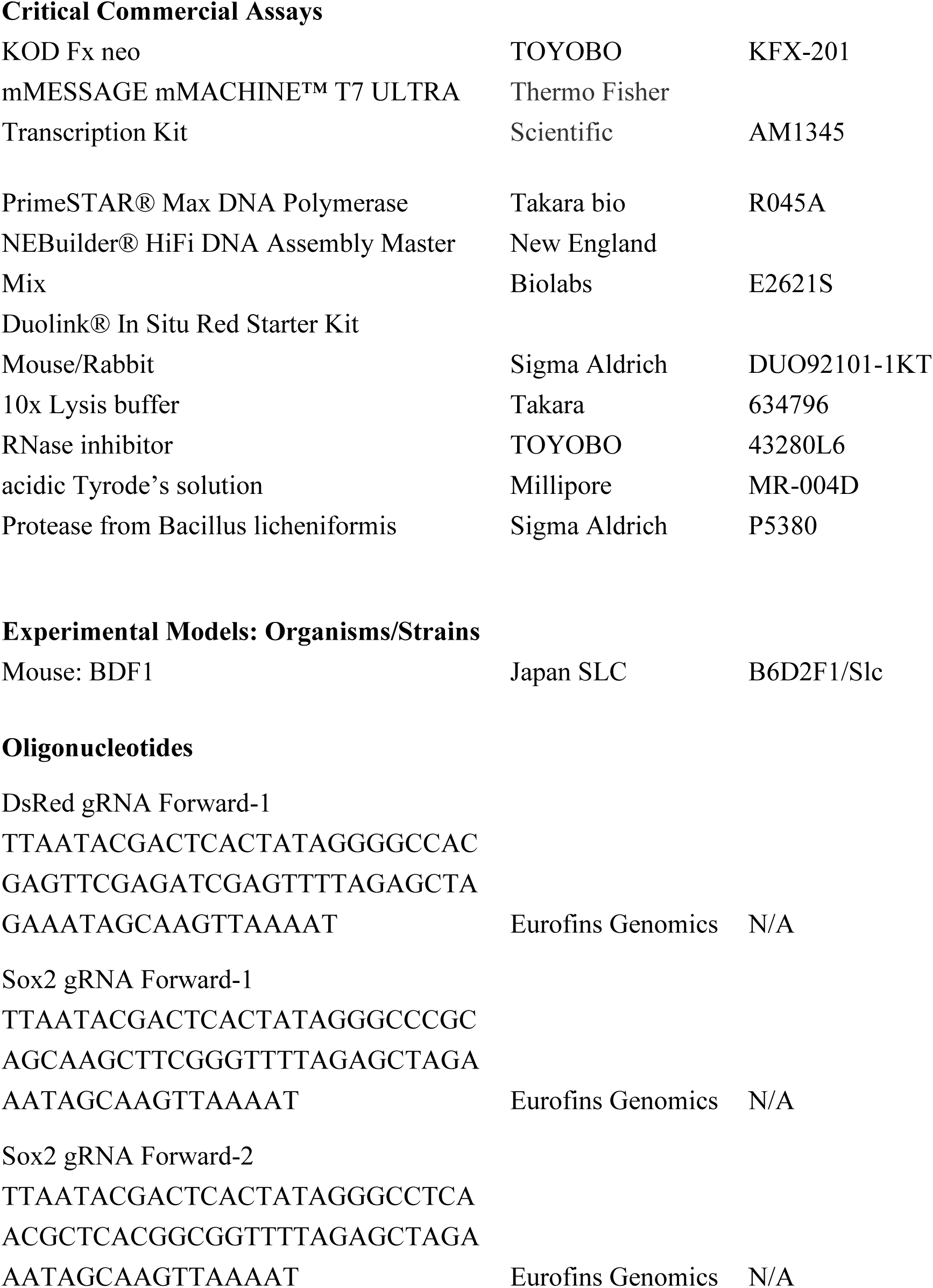
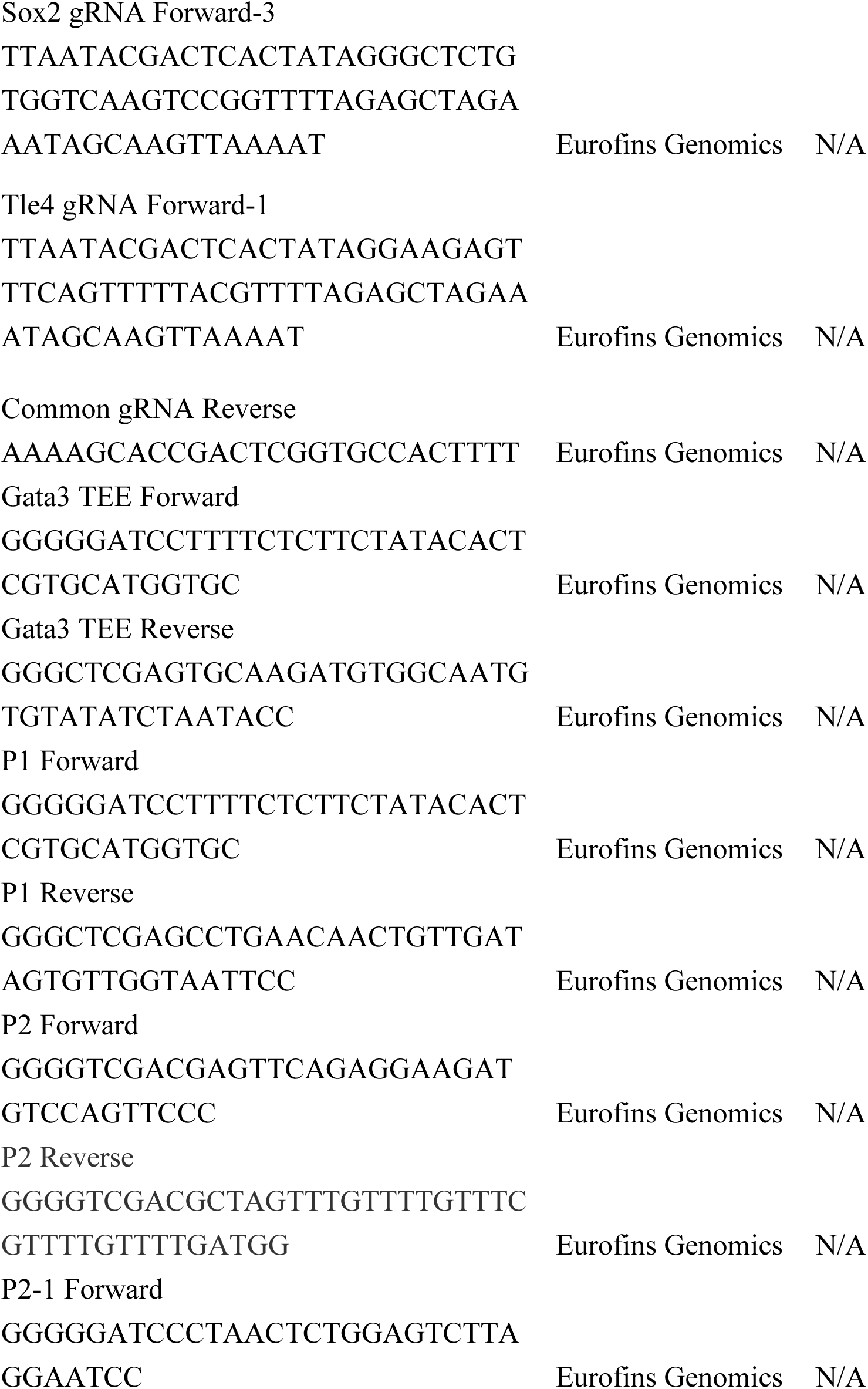
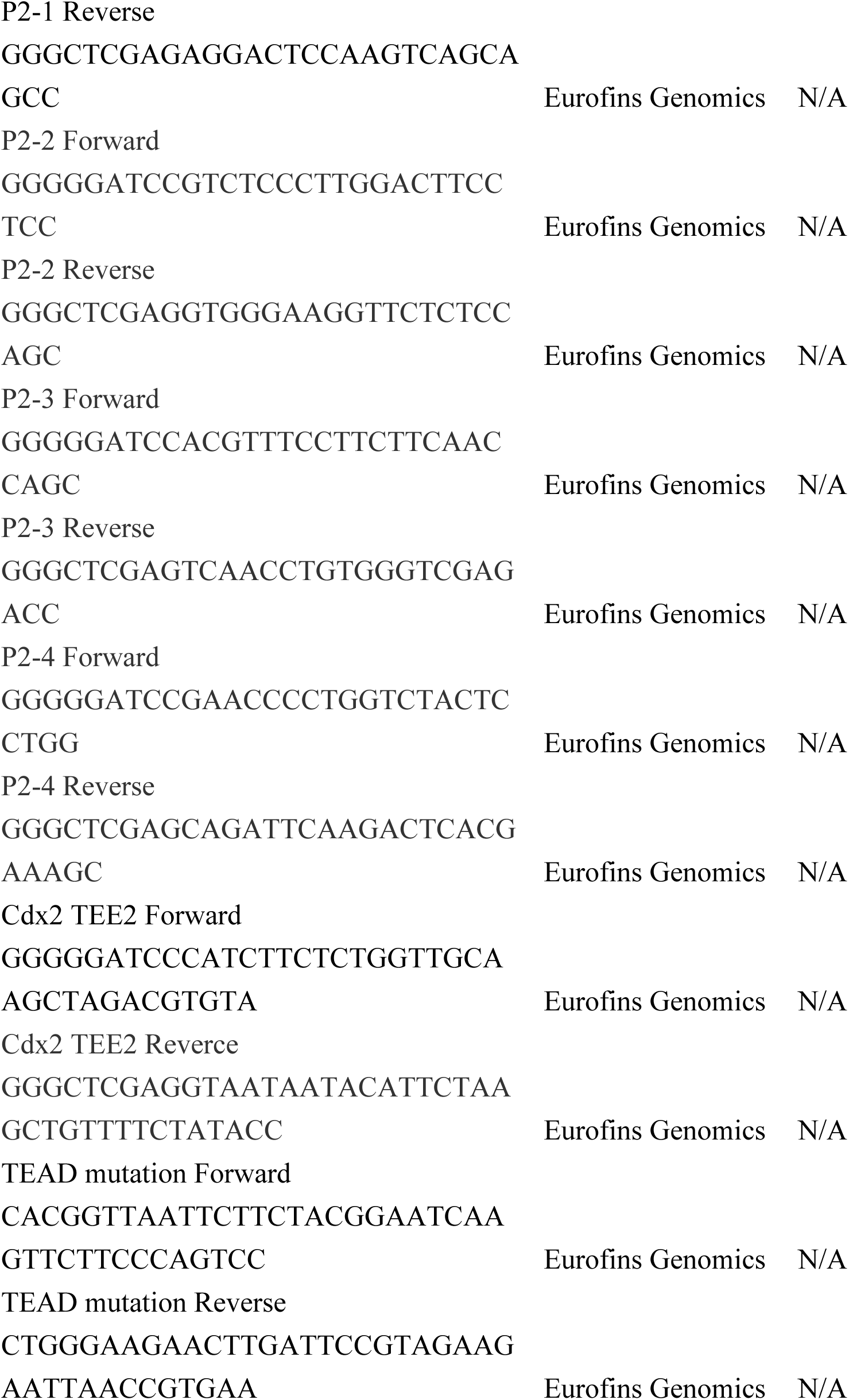
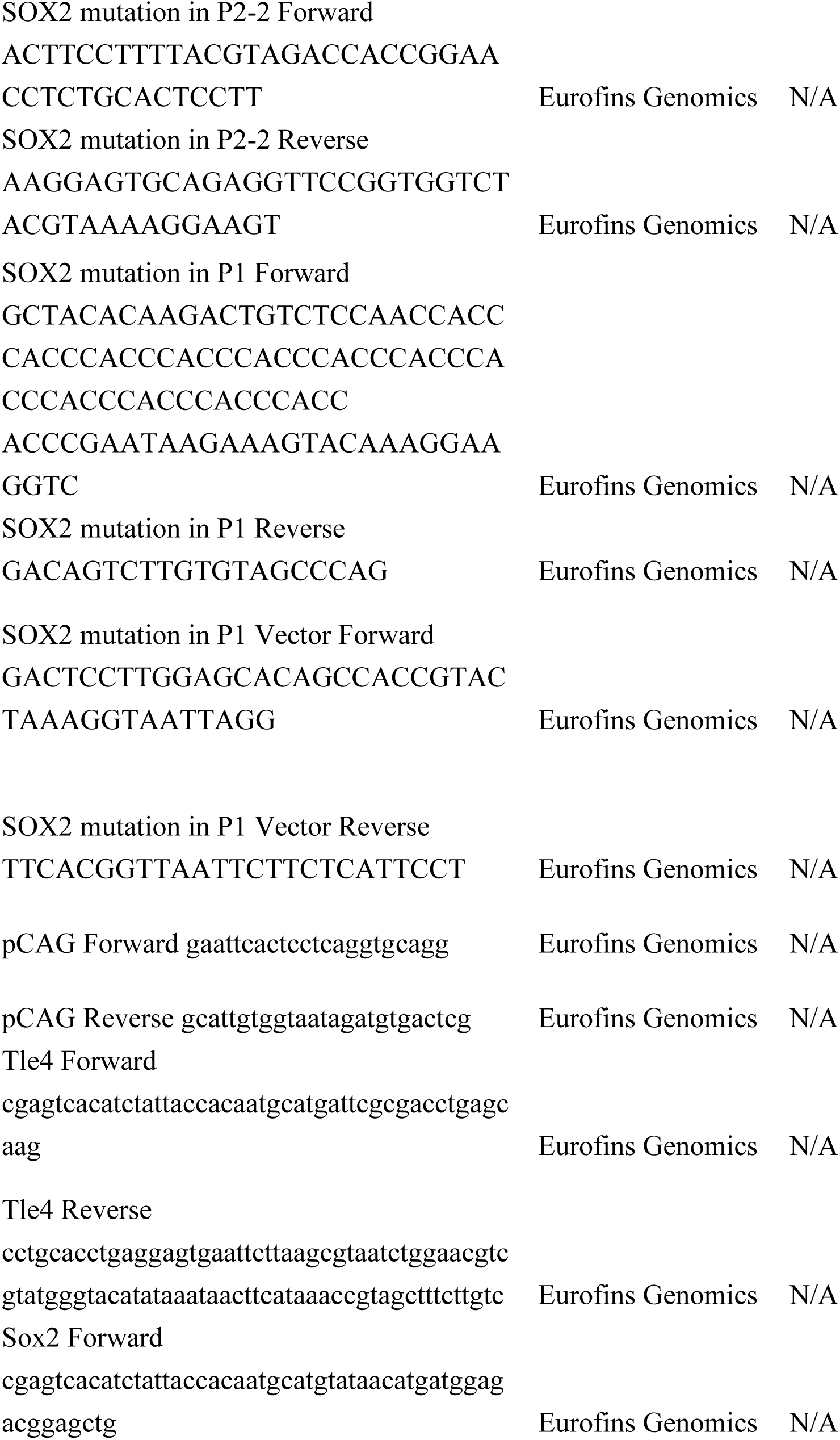
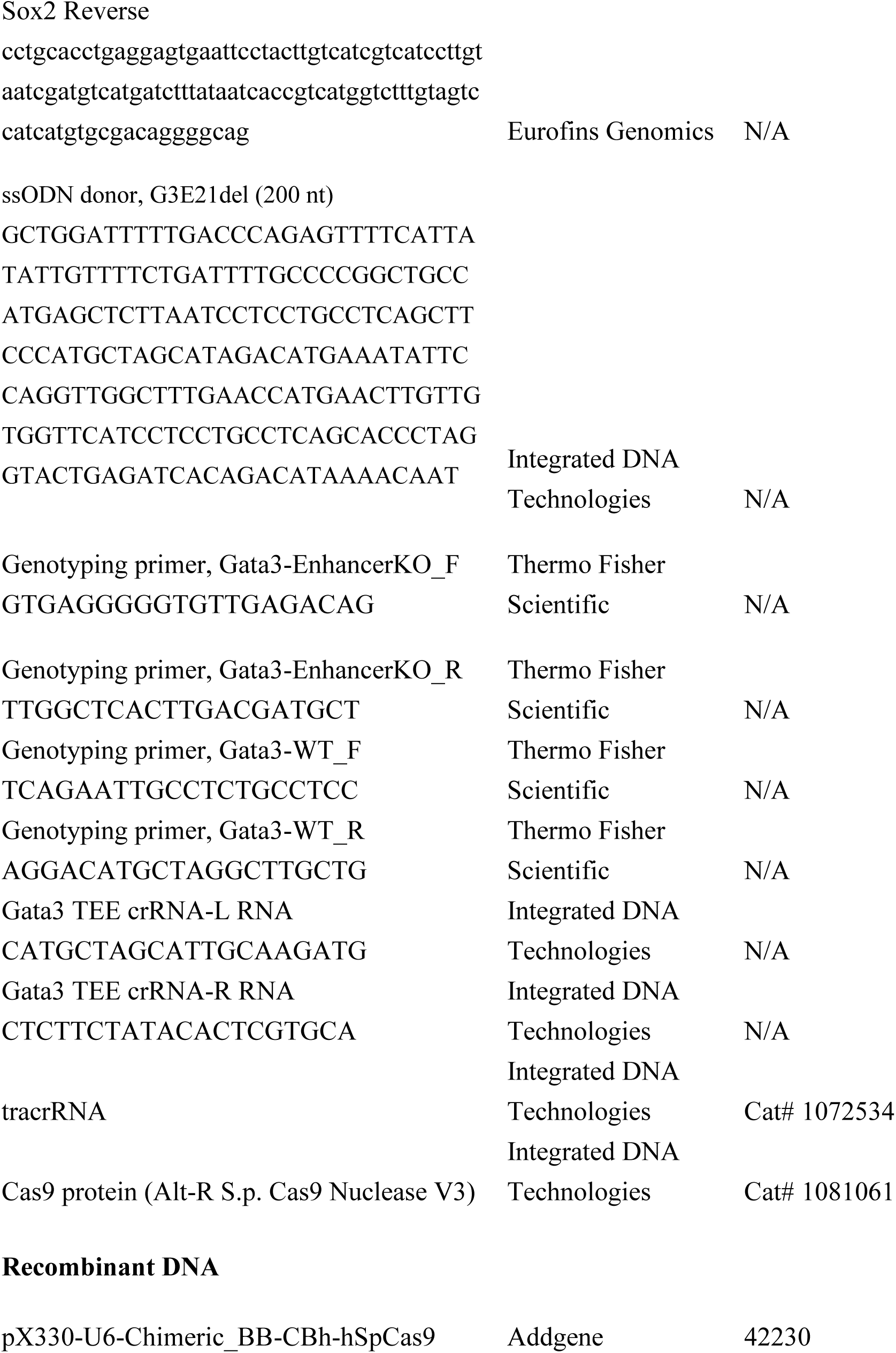
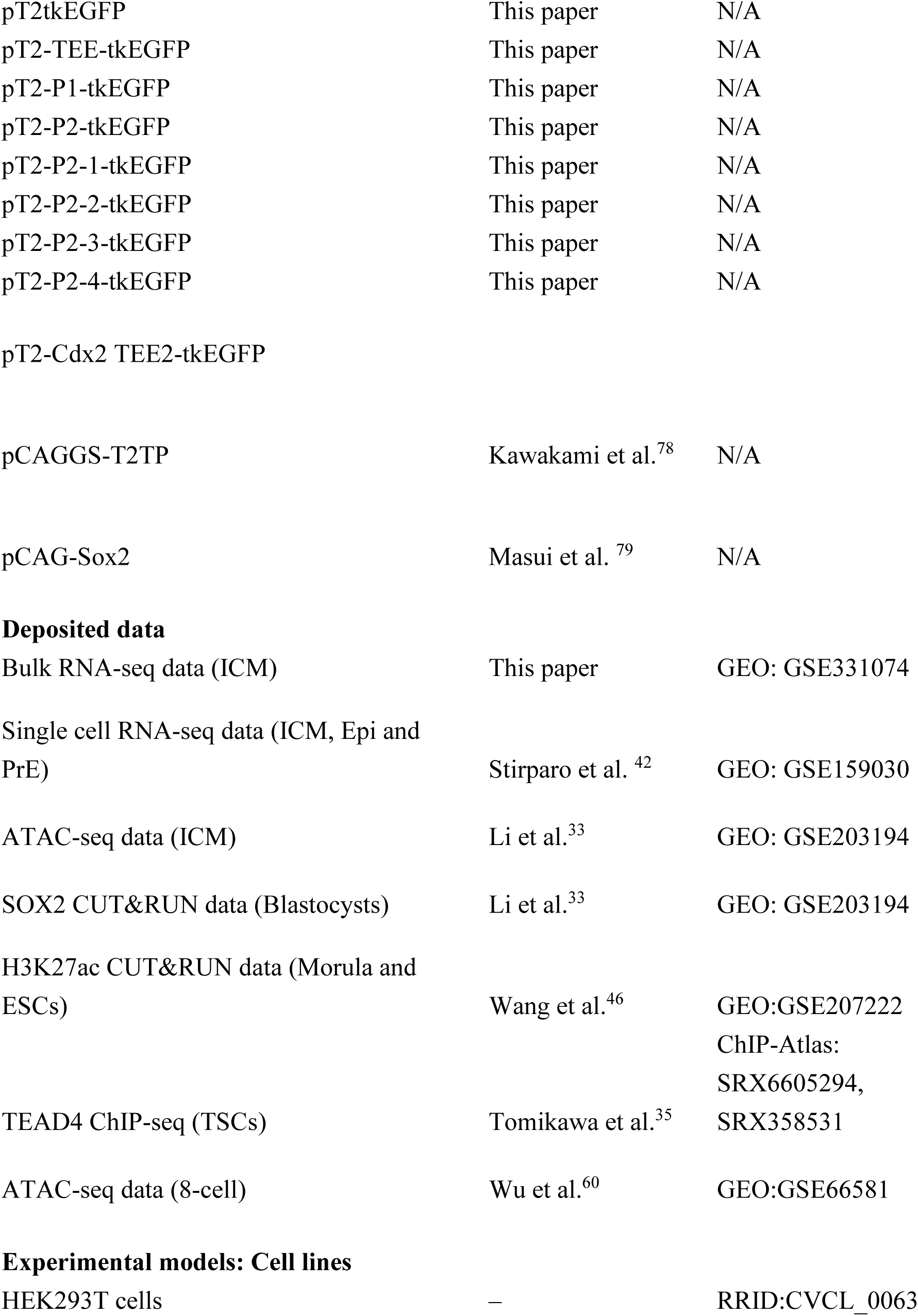
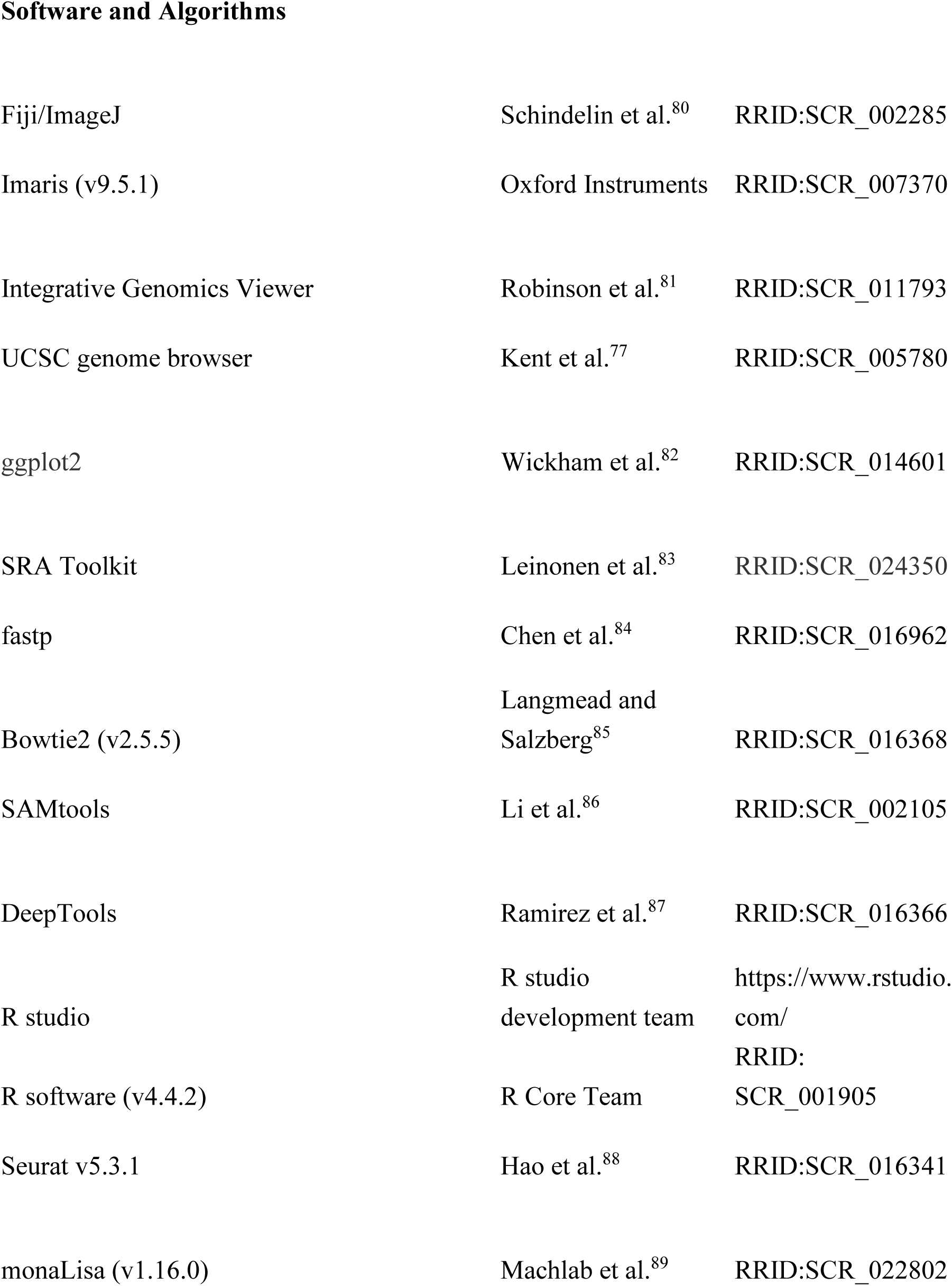
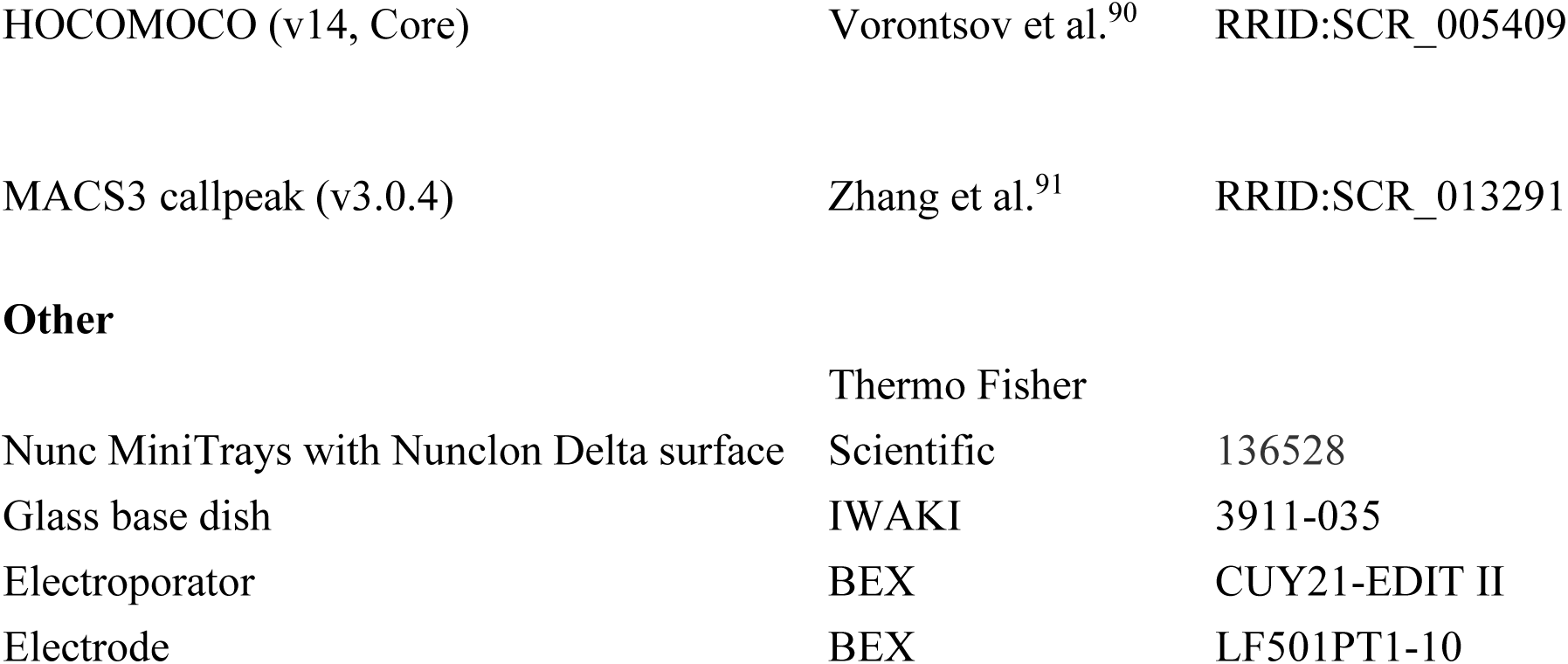

### EXPERIMENTAL MODEL AND STUDY PARTICIPANT DETAILS

#### Mice

B6D2F1/Jcl (hereafter referred to as BDF1) mice were obtained from CLEA Japan or generated in-house by crossing C57BL/6N Jcl (CLEA Japan) and DBA/2N Jcl (CLEA Japan). C57BL/6J mice (CLEA Japan) were also used in this study. Mice were housed in environmentally controlled rooms at the Animal Facilities of the Graduate School of Frontier Biosciences or the Graduate School of Medicine, the University of Osaka. All animal experiments were approved by the Animal Care and Use Committee of the Graduate School of Frontier Biosciences and the Graduate School of Medicine, the University of Osaka. All recombinant DNA experiments were approved by the Gene Modification Experiments Safety Committee of the University of Osaka.

### METHOD DETAILS

#### Embryo culture and manipulation

Preimplantation embryos were collected according to a standard protocol with minor modifications.^92^ Briefly, BDF1 female mice were superovulated by intraperitoneal injection of 10 IU pregnant mare serum gonadotropin (PMSG), followed 46–52 h later by injection of human chorionic gonadotropin (hCG), and then mated with BDF1 male mice. Zygotes were collected from the ampullae of the oviducts at E0.5 and treated with hyaluronidase to remove cumulus cells. Eight-cell-stage embryos were collected from the oviducts at E2.5. Embryos were cultured in 10 μL KSOMaa (hereafter referred to as KSOM) medium (ARK Resource) in individual wells of a 72-well MiniTray (Nunc, 136528), overlaid with 5 μL mineral oil, at 37°C in a humidified incubator with 5% CO_2_. E4.0 embryos used for the PLA were collected from the uterus by flushing with M2 medium.

#### Plasmid construction

The empty reporter plasmid used for enhancer reporter assays (*p*T2tkEGFP) was constructed by combining the Tol2 terminal repeats from pT2AL200R175^93^ with a reporter cassette consisting of the herpes simplex virus thymidine kinase minimal promoter, enhanced green fluorescent protein (EGFP), and a poly(A) signal.^94^ Reporter plasmids containing the *Gata3* TEE (*p*T2-TEE-tkEGFP) or the *Cdx2* TEE2 (*p*T2-*Cdx2* TEE2-tkEGFP) were generated by inserting the corresponding enhancer fragments into the *p*T2tkEGFP vector. Enhancer fragments were amplified by PCR using KOD Fx Neo DNA polymerase (Toyobo) and mouse kidney genomic DNA as the template. The *Gata3* TEE was amplified using the *Gata3* TEE Forward and Reverse primers (+56,048 to +61,466 bp relative to the TSS). The *Cdx2* TEE2 fragment was amplified using the *Cdx2* TEE2 Forward and Reverse primers (+6,428 to +9,996 bp relative to the TSS). Additional reporter plasmids containing TEE subfragments were generated using *p*T2-TEE-tkEGFP as the template. PCR amplification was performed using the indicated primer pairs. For *p*T2-TEE-tkEGFP, PCR products were digested with SalI and self-ligated. For the remaining constructs, inserts were digested with BamHI and XhoI and ligated into BamHI/XhoI-digested *p*T2-tkEGFP. The following primer pairs were used for plasmid construction: *p*T2-P1-tkEGFP, P1 Forward and P1 Reverse; *p*T2-P2-tkEGFP, P2 Forward and P2 Reverse; *p*T2-P2-1-tkEGFP, P2-1 Forward and P2-1 Reverse; *p*T2-P2-2-tkEGFP, P2-2 Forward and P2-2 Reverse; *p*T2-P2-3-tkEGFP, P2-3 Forward and P2-3 Reverse; and *p*T2-P2-4-tkEGFP, P2-4 Forward and P2-4 Reverse. Primer sequences are listed in the Key Resources Table.

To generate *p*CAG-*Tle4*–HA, a PCR product amplified from *p*CAGGS-T2TP using the pCAG Forward and Reverse primers and a PCR product amplified from *Tle4* cDNA (DNABook Mouse Transcription Factors, DNAFORM) using the *Tle4* Forward and Reverse primers were assembled using the NEBuilder HiFi DNA Assembly System (New England Biolabs) according to the manufacturer’s instructions. To generate *p*CAG-*Sox2*–FLAG×3, a PCR product amplified from *p*CAGGS-T2TP using the pCAG Forward and Reverse primers and a PCR product amplified from a plasmid containing mouse *Sox2* cDNA (*p*CAG-*Sox2*, kindly provided by Dr. Hitoshi Niwa)^79^ using the *Sox2* Forward and Reverse primers were assembled using the NEBuilder HiFi DNA Assembly System (New England Biolabs) according to the manufacturer’s instructions. Primer sequences are listed in the Key Resources Table.

#### Mutagenesis

Mutant enhancer reporter plasmids were generated by site-directed mutagenesis using PrimeSTAR Max DNA Polymerase (Takara Bio) or the NEBuilder HiFi DNA Assembly System (New England Biolabs) according to the manufacturers’ instructions. The TEAD-binding motif within the P2-2 region (5′-TCTCATTCCTCA-3′) was mutated to 5′-TCTacggaaTCA-3′. The SOX2-binding motif within the P2-2 region (5′-AGAAACAATTAAC-3′) was mutated to 5′-AGAccaccggAAC-3′. The 12 SOX2-binding motifs within the P1 region (5′-CAAAACAAACAAACAAACAAACAAACAAACAAACAAACAAACAAACAAAAAACAAAGAA-3′) was mutated to 5′-CAAccacccacccacccacccacccacccacccacccacccacccacccAccacccGAA-3′. Lowercase letters indicate the mutated nucleotides. Each mutation was introduced using the following primer pairs: mut TEAD, TEAD mutation Forward and Reverse; mut SOX2 in P1, SOX2 mutation in P1 Forward and Reverse; and mut SOX2 in P2-2, SOX2 mutation in P2-2 Forward and Reverse. Primer sequences are listed in the Key Resources Table.

#### Knockout embryo production

*Sox2* and *Tle4* knockout embryos were generated by CRISPR/Cas9-mediated genome editing of zygotes, as previously described.^34,95^ Each sgRNA template was generated by PCR amplification of pX330-U6-Chimeric_BB-CBh-hSpCas9 using the following primer pairs: DsRed gRNA Forward-1 and Common gRNA Reverse; *Sox2* gRNA Forward-1 and Common gRNA Reverse; *Sox2* gRNA Forward-2 and Common gRNA Reverse; *Sox2* gRNA Forward-3 and Common gRNA Reverse; and *Tle4* gRNA Forward-1 and Common gRNA Reverse. Primer sequences are listed in the Key Resources Table. PCR products were purified by ethanol precipitation, washed with 70% ethanol, and air-dried before resuspension. gRNAs were synthesized by in vitro transcription from the purified PCR products using the MEGAshortscript T7 Transcription Kit (Thermo Fisher Scientific) according to the manufacturer’s instructions. The synthesized gRNAs were purified by phenol–chloroform– isoamyl alcohol extraction followed by isopropanol precipitation and resuspended in RNase-free water. Cas9 protein (Integrated DNA Technologies) and gene specific gRNAs were introduced into the zygotes by electroporation. Three gRNAs were used to target *Sox2*, whereas a single gRNA was used to target *DsRed* (control) or *Tle4*.

#### Transgenic embryo production

Transgenic embryos for enhancer reporter assays were generated by pronuclear microinjection of the *p*T2tkEGFP plasmid containing the indicated enhancer fragments together with *Tol2* transposase mRNA. Microinjection was performed using a micromanipulator mounted on an inverted microscope as previously described.^92^ Plasmid DNA (10 ng/μL) and *Tol2* transposase mRNA (100 ng/μL) were mixed in nuclease-free water and injected into the pronuclei of one-cell-stage zygotes^78^. *Tol2* transposase mRNA was synthesized by *in vitro* transcription from a Tol2 transposase expression plasmid *p*CAGGS-T2TP ^96^ by using the mMESSAGE mMACHINE T7 Ultra Kit (Thermo Fisher Scientific) according to the manufacturer’s instructions. The mRNA was purified by phenol– chloroform–isoamyl alcohol extraction followed by isopropanol precipitation.

#### Generation of the *Gata3^ΔTEE^* mouse line

The *Gata3* TEE-deleted mouse line (*Gata3^ΔTEE^*) was generated by electroporation of Cas9 ribonucleoprotein (RNP) complexes into C57BL/6J zygotes. Two crRNAs targeting sequences flanking the TEE (*Gata3* TEE crRNA-L, CATGCTAGCATTGCAAGATG; and *Gata3* TEE crRNA-R, CTCTTCTATACACTCGTGCA), tracrRNA, and Cas9 protein (Alt-R S.p. Cas9 Nuclease V3) were purchased from Integrated DNA Technologies. Each crRNA was annealed with tracrRNA to generate a crRNA:tracrRNA duplex (gRNA). A 200-nt single-stranded oligodeoxynucleotide (ssODN) spanning the expected deletion junction was used as the repair template. The crRNA and ssODN sequences are listed in the Key Resources Table.

Cryopreserved C57BL/6J zygotes produced by in vitro fertilization using oocytes from superovulated C57BL/6J females and sperm from C57BL/6J males were used for electroporation. The zygotes had been cryopreserved by vitrification until use. Electroporation was performed as previously described.^95^ Opti-MEM I (Thermo Fisher Scientific, 11058-021) supplemented with 0.1% polyvinyl alcohol (Sigma-Aldrich, P8136) and sterilized by filtration was used as the electroporation solution. Thawed zygotes were washed twice with the electroporation solution and once with the same solution containing the two gRNAs (100 ng/μL in total), Cas9 protein (100 ng/μL), and ssODN (300 ng/μL). The zygotes were aligned within the 1-mm gap of an LF501PT1-10 platinum plate electrode (BEX) filled with 10 μL of the same ribonucleoprotein-containing solution. Electroporation was performed using a CUY21EDIT II electroporator (BEX) with five pulses at 30 V (3 ms on, 97 ms off). After electroporation, zygotes were washed twice with modified Whitten’s medium (mWM)^97^ and cultured at 37°C under 5% CO_2_. Embryos that developed to the two-cell stage were transferred into the oviducts of pseudopregnant recipient females.

Founder mice were identified by genomic PCR and crossed with C57BL/6J mice to establish the line. Genotyping was performed by genomic PCR using TaKaRa Ex Premier DNA Polymerase (Takara Bio) in 25-μL reaction volumes for 30 cycles with an annealing temperature of 60°C and an extension time of 45 s; all other conditions followed the manufacturer’s instructions. The *Gata3^ΔTEE^*deleted allele was detected as a 721-bp PCR product using the Gata3-EnhancerKO_F and Gata3-EnhancerKO_R primers, whereas the wild-type allele was detected as a 1,009-bp PCR product using the Gata3-WT_F and Gata3-WT_R primers. Primer sequences are listed in the Key Resources Table. The deleted allele was further verified by Sanger sequencing, which confirmed deletion of the entire TEE (Figure S4F). *Gata3^ΔTEE^* mice were maintained on a C57BL/6J genetic background.

#### Inhibitor treatments

For inhibitor treatments, embryos were cultured in KSOM medium containing either 10 μM TRULI (MedChemExpress), a LATS inhibitor,^28^ or 3 mM sodium butyrate (Selleck), an HDAC inhibitor.^98,99^ Control embryos were cultured in KSOM containing the corresponding concentration of DMSO for TRULI treatment or in KSOM alone for sodium butyrate treatment.

#### Immunofluorescent staining and confocal image acquisition

Embryos were subjected to immunofluorescence staining using a 72-well MiniTray (Nunc, 136528), as previously described.^21^ Briefly, embryos were fixed in 4% paraformaldehyde in Dulbecco’s phosphate-buffered saline (PBS) for 15 min at room temperature, and then permeabilized by washing twice with 0.1% Triton X-100 in PBS (PBST) at room temperature for 1 min each. Embryos were blocked in 2% donkey serum in PBST for 5 min and then incubated overnight at 4°C with primary antibodies diluted 1:100 in blocking solution. After washing twice with PBST for 1 min each, embryos were incubated for at least 1 h at room temperature with secondary antibodies diluted 1:1000 in PBST containing 1 μg/mL Hoechst 33342 (Dojindo). Primary and secondary antibodies are listed in the Key Resources Table. For image acquisition, embryos were placed in a small drop of PBS on glass-bottomed dishs (IWAKI). Confocal images were acquired using either a spinning-disk confocal microscope consisting of a Nikon Ti2 microscope, a Dragonfly imaging system (Oxford Instruments), and a Zyla sCMOS camera (Oxford Instruments), or a Nikon A1 point-scanning confocal microscope. Images were analyzed using Imaris (Bitplane) or Fiji/ImageJ (National Institutes of Health).

#### Proximity ligation assay

The proximity ligation assay (PLA) was performed using the Duolink In Situ Red Starter Kit (Sigma-Aldrich), as previously described.^100^ E4.0 embryos were fixed in 4% paraformaldehyde for 10 min at room temperature, permeabilized with 0.1% Triton X-100 in PBS (PBST), and blocked with Duolink Blocking Solution for 1 h at 37°C. Primary antibodies for PLA (rabbit anti-SOX2 and mouse anti-TLE4) and immunofluorescence (goat anti-SOX2) were diluted 1:100 in Duolink Antibody Diluent and incubated with embryos overnight at 4°C. After washing with 1× Wash Buffer A, embryos were incubated with PLA probes (PLUS and MINUS) for 1 h at 37°C. Ligation was performed using Duolink Ligation Solution for 30 min at 37°C, followed by amplification with Duolink Amplification Solution Red for 120 min at 37°C. Embryos were washed with Wash Buffer B and subsequently processed for immunofluorescence staining with secondary antibodies and Hoechst 33342, as described above.

#### Cell culture

100 × 20 mm dishes (Thermo Fisher Scientific) were coated with gelatin (Cellmatrix Type I-P) (Nitta Gelatin) prepared in 1 mM HCl (pH 3.0) and incubated for 30–60 min at room temperature. HEK293T cells were maintained in Dulbecco’s Modified Eagle Medium (DMEM) (FUJIFILM) supplemented with 10% fetal bovine serum (FBS) (Sigma Aldrich) at 37°C under 5% CO₂.

#### Co-immunoprecipitation and immunoblotting

One day before transfection, 4.4 × 10⁶ HEK293T cells were plated onto each gelatin-coated 100 × 20 mm dish (Thermo Fisher Scientific). Twenty-four hours after plating, 8 μg of each plasmid DNA (*p*CAG-Sox2–FLAGx3 and pCAG-Tle4–HA) was transfected using 20 μL of Lipofectamine 2000 (Thermo Fisher Scientific) according to the manufacturer’s instructions. After 44 h post-transfection, the cells were washed with PBS and treated with a cross-linking solution (2 mM dithio-bis (succinimidyl propionate) (Sigma-Aldrich) in PBS) for 30 min. The cross-linking reaction was stopped by incubation with 20 mM Tris-HCl (pH 7.5) for 15 min at room temperature. The cells were then lysed in 1 mL of RIPA buffer (50 mM Tris-HCl, pH 7.6; 150 mM NaCl; 1 mM EDTA; 10% glycerol; 1% Triton X-100; 0.5% sodium deoxycholate; 0.1% SDS; cOmplete™ Protease Inhibitor Cocktail) for 30 min and harvested using a cell scraper. The lysates were sonicated twice for 30 s with a 30 s interval and centrifuged at 13,000 rpm. The supernatants were used as the cell lysates.

Forty microliters of Dynabeads™ Protein G (Invitrogen) per sample were incubated with 2 μg of anti-HA [M180-3] antibody or 10 μg of anti-FLAG [M2] antibody diluted in 200 μL of PBST (PBS containing 0.1% Tween 20) for more than 1 h at room temperature. After incubation, the beads were washed once with 0.1% Tween 20 in PBS. The antibody-conjugated beads were incubated with 300 μL of cell lysate overnight at 4°C. The beads were washed three times with wash buffer (50 mM Tris-HCl, pH 7.6; 150 mM NaCl; 1 mM EDTA; 0.1% NP-40; cOmplete™ Protease Inhibitor Cocktail [Roche]), resuspended in 30 μL of wash buffer, and mixed with 30 μL of 4× SDS sample buffer, followed by heating at 95°C for 5 min. For the 5% input sample, 20 μL of lysate was mixed with 20 μL of 4× SDS sample buffer. After heating, the beads were separated using a magnetic rack, and the supernatant was collected.

For immunoblotting, 5 μL of the input sample, 10 μL of the immunoprecipitated sample, and 8 μL of Precision Plus Protein™ Dual Color Standards (Bio Rad) were separated on a 12.5% polyacrylamide gel by SDS-polyacrylamide gel electrophoresis. Proteins were transferred onto Whatman® Protran® nitrocellulose membranes (Cytiva) and stored in TBST (25 mM Tris-HCl, 2.7 mM KCl, 140 mM NaCl, and 0.1% Tween 20) at 4°C. The membranes were blocked with Blocking One (Nacalai Tesque) for 2 h at room temperature with gentle agitation. After washing with TBST, the membranes were incubated with primary antibodies (anti-HA [3F-10] or anti-FLAG[M2]) diluted in Can Get Signal® Immunoreaction Enhancer Solution 1 (TOYOBO) for 2 h at room temperature. The membranes were then washed with TBST and incubated with HRP-conjugated secondary antibodies (anti-rat or anti-mouse) diluted in Can Get Signal® Solution 2 for 2 h at room temperature. After washing with TBST, signals were detected using signals were detected using a ChemiDoc MP Imaging System (Bio-Rad).

#### Bulk RNA-seq

ICMs from *Sox2* KO embryos or control *DsRed* KO embryos generated by genome editing of zygotes were used for Bulk RNA-seq. At the 8-cell stage, the zona pellucida was removed using acidic Tyrode’s solution (Millipore), and the embryos were subsequently treated with a LATS inhibitor for 12 h beginning at the mid-blastocyst stage. Treated embryos were washed in M2 medium and incubated in M2 medium supplemented with 0.5 M EDTA (pH 8.0) and 5 μg/μL protease from Bacillus licheniformis (Sigma-Aldrich) for TE isolation. After removal of the TE layer, each isolated ICMs were washed with PBS and individually lysed using lysis buffer (Takara) supplemented with RNase inhibitor (TOYOBO). Each isolated ICM was frozen in liquid nitrogen until use. Library preparation was performed using the SMART-Seq mRNA HT LP Kit (Takara) according to the manufacturer’s instructions. Sequencing was performed on a NovaSeq X Plus platform (Illumina) in 151-nt single-end mode. After adapter trimming using Trimmomatic version 0.39, the generated reads were mapped to the mouse reference genome (mm10) using HISAT2 version 2.1.0. Differentially expressed genes (DEGs) were detected using DESeq2 (v1.50.2) with the Wald test. The adjusted *p*-value threshold was set to 0.1. Gene Ontology enrichment analysis was performed using the R package *clusterProfiler* (v4.18.4) together with the annotation database *org.Mm.eg.db* (v3.22.0). For visualization, the top five enriched terms are shown.

### QUANTIFICATION AND STATISTICAL ANALYSIS

#### Visualization of published genomics datasets

For datasets for which BigWig files were directly available from the NCBI Gene Expression Omnibus (GEO) or ChIP-Atlas, the files were downloaded and loaded into the Integrative Genomics Viewer (IGV)^81^ without further processing. For datasets for which BigWig files were not available from NCBI GEO, the publicly available genomic datasets were re-analyzed as follows. Raw sequencing data (SRR files) obtained from NCBI GEO were converted to FASTQ format using fastq-dump from the SRA Toolkit, followed by adapter trimming and quality filtering using fastp^84^. Processed reads were aligned to the mouse reference genome (mm9) using Bowtie2^85^ with the default parameters. BAM files were generated, sorted, and indexed using SAMtools^86^. To enable comparisons across samples, the BAM files were normalized to counts per million and converted to BigWig format with a bin size of 10 bp using deepTools^87^ bamCoverage. All datasets were visualized using the mm9 genome assembly, and the signal intensities at the genomic regions of interest were compared. The public datasets used in this study are listed in the Key Resources Table.

#### ATAC-seq peak clustering

ATAC-seq and CUT&RUN datasets were obtained from the GEO (GSE203194). ATAC-seq datasets from E3.5 control embryos were used for peak identification. Reads were aligned to the mm9 reference genome using Bowtie2 (v2.5.5) with the --very-sensitive option. After alignment, unpaired or discordant reads were excluded using the corresponding Bowtie2 options. After BAM files from replicate samples were merged, peaks were called using MACS3 callpeak (v3.0.4)^91^. Peaks with a *q* value ≤ 0.05 were retained for downstream analysis.

Signal values were extracted from the BigWig files provided as processed data in GEO (GSM7211420–GSM7211425). For each peak, the maximum signal value within the overlapping genomic region was used. Peaks were clustered based on accessibility patterns across four samples: E3.5 control, E4.5 control, E3.5 *Sox2* KO, and E4.5 *Sox2* KO. Clustering was performed using k-means, with the number of clusters set to four based on an elbow plot. Log_2_ fold changes relative to E3.5 control were calculated and visualized as a heatmap.

Candidate peaks of interest were selected based on ATAC-seq and SOX2 CUT&RUN signals. SOX2 CUT&RUN signal levels at each peak were quantified using the same procedure as for ATAC-seq. For both datasets, peaks within the top 30% of signal values were defined as high-signal peaks. Peaks classified as high-signal peaks in both datasets were selected for subsequent analysis.

#### Analysis of the nearest TE genes for candidate peaks

To calculate the Jaccard coefficients, the nearest gene was assigned to each peak based on the distance to the TSS, and gene lists were generated for each cluster. TE marker genes reported by Proks et al.^52^ were classified into E3.5-specific, E3.5–E4.5 common, and E4.5-specific gene sets. Jaccard coefficients were then calculated by comparing each nearest-gene list with these marker gene sets.

#### Motif enrichment for candidate peaks

Motif enrichment analysis was performed using DNA sequences corresponding to the selected peaks and the R package *monaLisa* (v1.16.0)^89^. HOCOMOCO (v14, Core)^90^ was used as the motif database. Sequence-based motif-specific bias was normalized using genome-wide background information by setting the background = "genome" option.

#### Analysis of published scRNA-seq data

The scRNA-seq dataset from a previous study (GSE159030)^42^ was reanalyzed. Ensembl gene IDs were mapped to the corresponding gene symbols. Downstream analysis was performed in the R software environment using Seurat^88^, ggplot2^82^, and patchwork.

#### Statistical analysis

Statistical analyses were performed using R. The statistical methods and numbers of samples (N, number of embryos; n, number of cells) are described in the figures or figure legends.

