## Supplemental Figures S1-S6 for "SOX2 terminates trophectoderm competence in inner cell mass by closing trophectoderm enhancers"

Supplemental information

Document S1. Figures S1–S6

### Figure S1, related to Figure 2

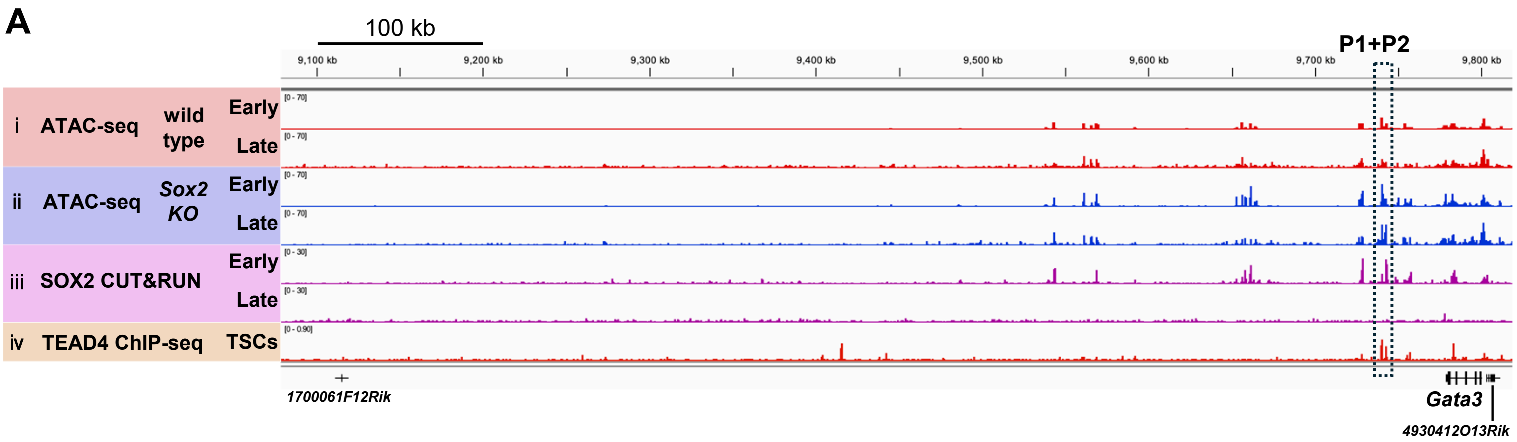


**Figure S1. Chromatin accessibility and transcription factor binding sites between *Gata3* and neighboring genes, related to Figure 2.**

(A) Reduced-scale IGV snapshot corresponding to Figure 2A. The dashed box indicates P1+P2. Early, ICM of early blastocysts; Late, ICM of late blastocysts; TSCs, trophoblast stem cells.

### Figure S2, related to Figure 3

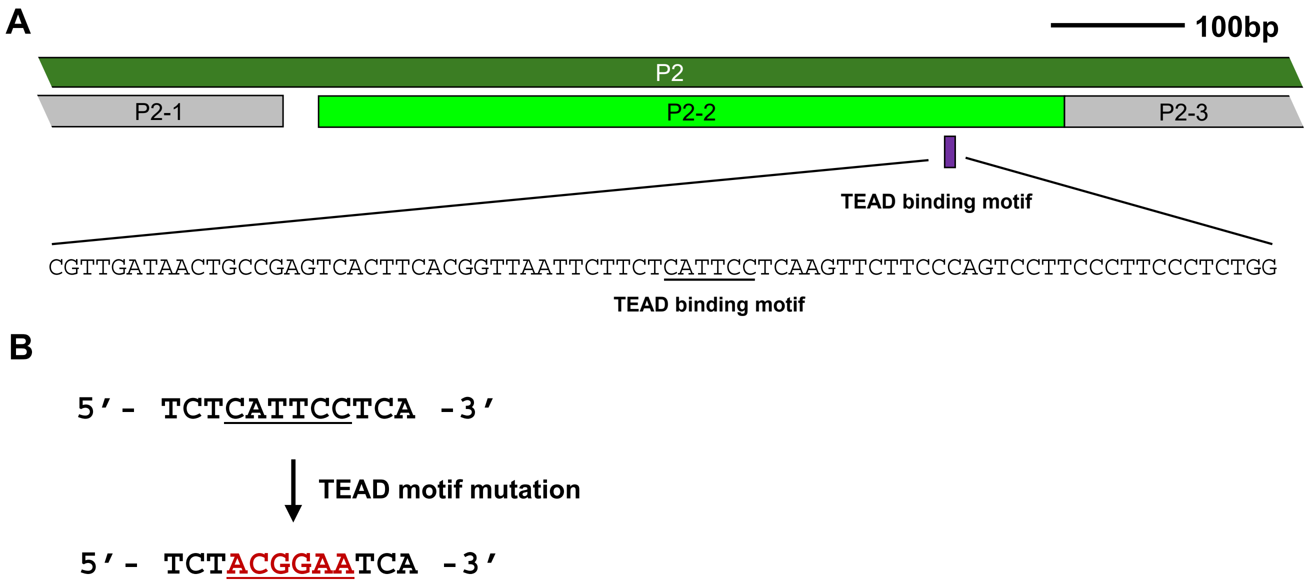


**Figure S2. TEAD-binding motif within P2-2, related to Figure 3.**
(A) Schematic showing the position of the TEAD-binding motif within P2-2 and the surrounding DNA sequence (bottom). Scale bar, 100 bp.
(B) Schematic showing the TEAD motif mutation introduced by base substitutions (A↔C and G↔T).

### Figure S3, related to Figure 4

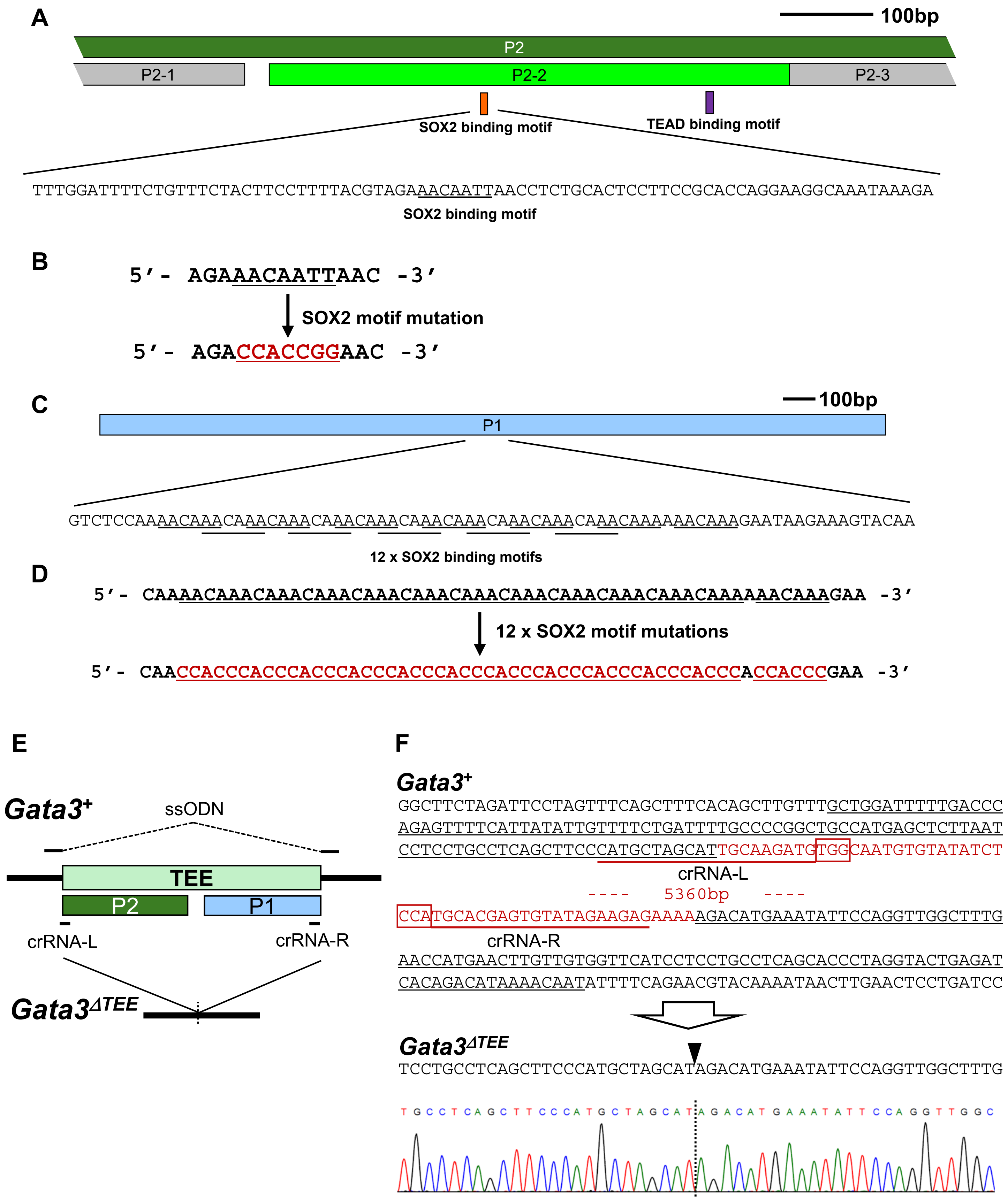

**Figure S3. SOX2-binding motifs within the TEE, related to Figure 4.**
(A) Schematic showing the position of the SOX2-binding motif within P2-2 and the surrounding DNA sequence (bottom). Scale bar, 100 bp.
(B) Schematic showing the SOX2 motif mutation introduced by base substitutions (A↔C and G↔T).
(C) Schematic showing the positions of SOX2-binding sites within P1 and the DNA sequences surrounding the SOX2-binding motifs (bottom). Scale bar, 100 bp.
(D) Schematic showing the mutations introduced into the 12 SOX2-binding motifs within P1.
(E) Schematic of the strategy used to generate the $Gata3^{\Delta TEE}$ mouse line. The positions of the crRNAs and ssODN used for genome editing are indicated.
(F) Sequences of the wild-type ($Gata3^{+}$) and $Gata3^{\Delta TEE}$ alleles surrounding the deletion sites. In the $Gata3^{+}$ sequence, the TEE sequence is shown in red. Thin underlines indicate the position of the ssODN, thick red underlines indicate the crRNAs, and red boxes indicate the PAM sequences. The DNA sequencing electropherogram confirms the expected deletion in the $Gata3^{\Delta TEE}$ allele.

### Figure S4, related to Figure 5

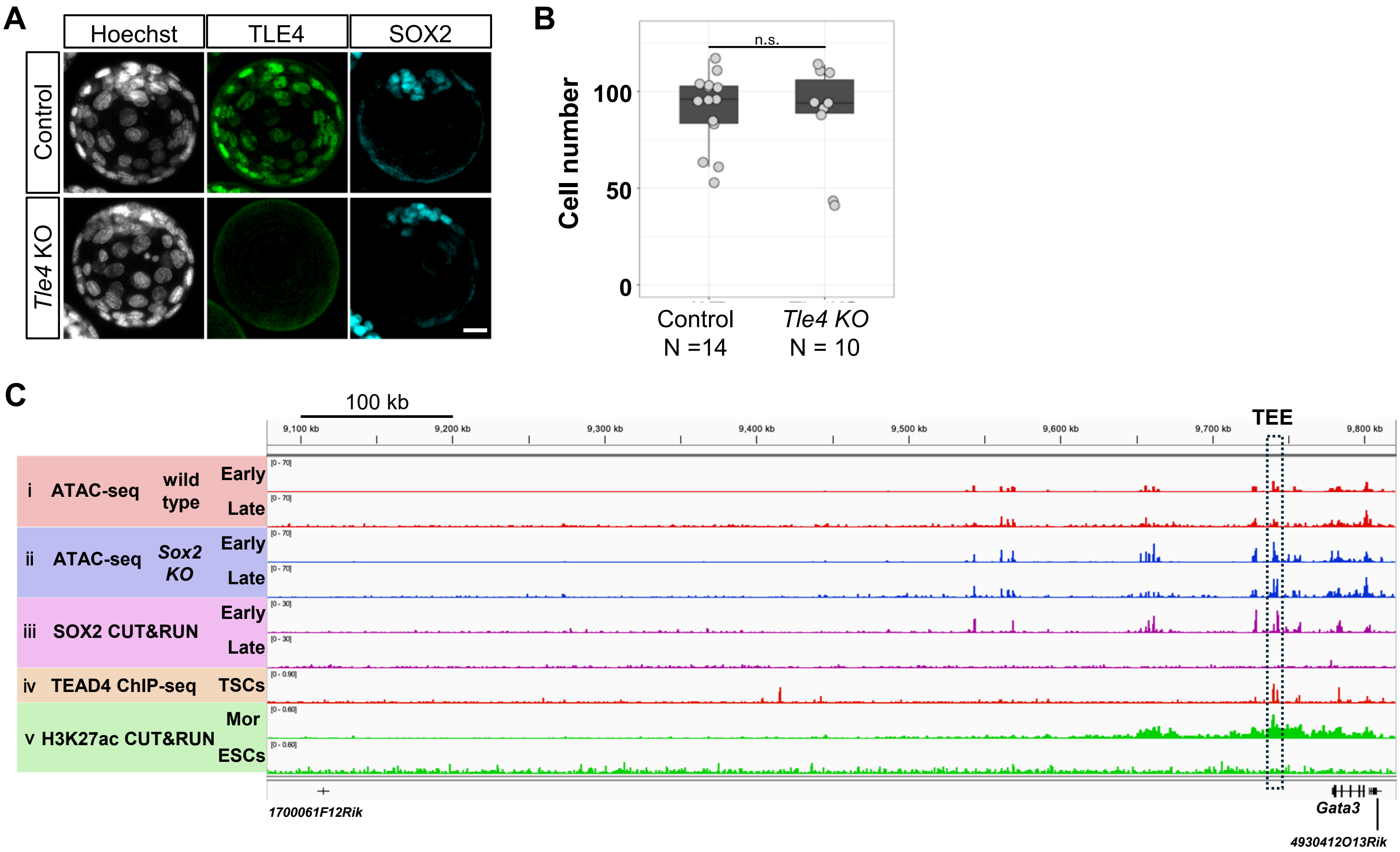


**Figure S4. Characterization of *Tle4* KO embryos, related to Figure 5.**
(A) Representative immunofluorescence images of TLE4 and SOX2 expression in control and *Tle4 KO* embryos. Images are shown as z-stack projections. Scale bar, 20 µm.
(B) Total cell numbers in the embryos shown in (A). Each dot represents an individual embryo. n.s., not significant (two-tailed unpaired Student's *t*-test).
(C) IGV snapshots of the genomic region surrounding the *Gata3* locus showing (i) ATAC-seq signals in ICMs of wild-type early and late blastocysts, (ii) ATAC-seq signals in ICMs of *Sox2 KO* early and late blastocysts, (iii) SOX2 CUT&RUN signals in early and late blastocysts, (iv) TEAD4 ChIP-seq signals in TSCs, and (v) H3K27ac CUT&RUN signals in morulae and ESCs. The dashed box indicates the TEE. Early, ICMs of early blastocysts; Late, ICMs of late blastocysts; TSCs, trophoblast stem cells; Mor, morulae; ESCs, embryonic stem cells.

### Figure S5, related to Figure 6

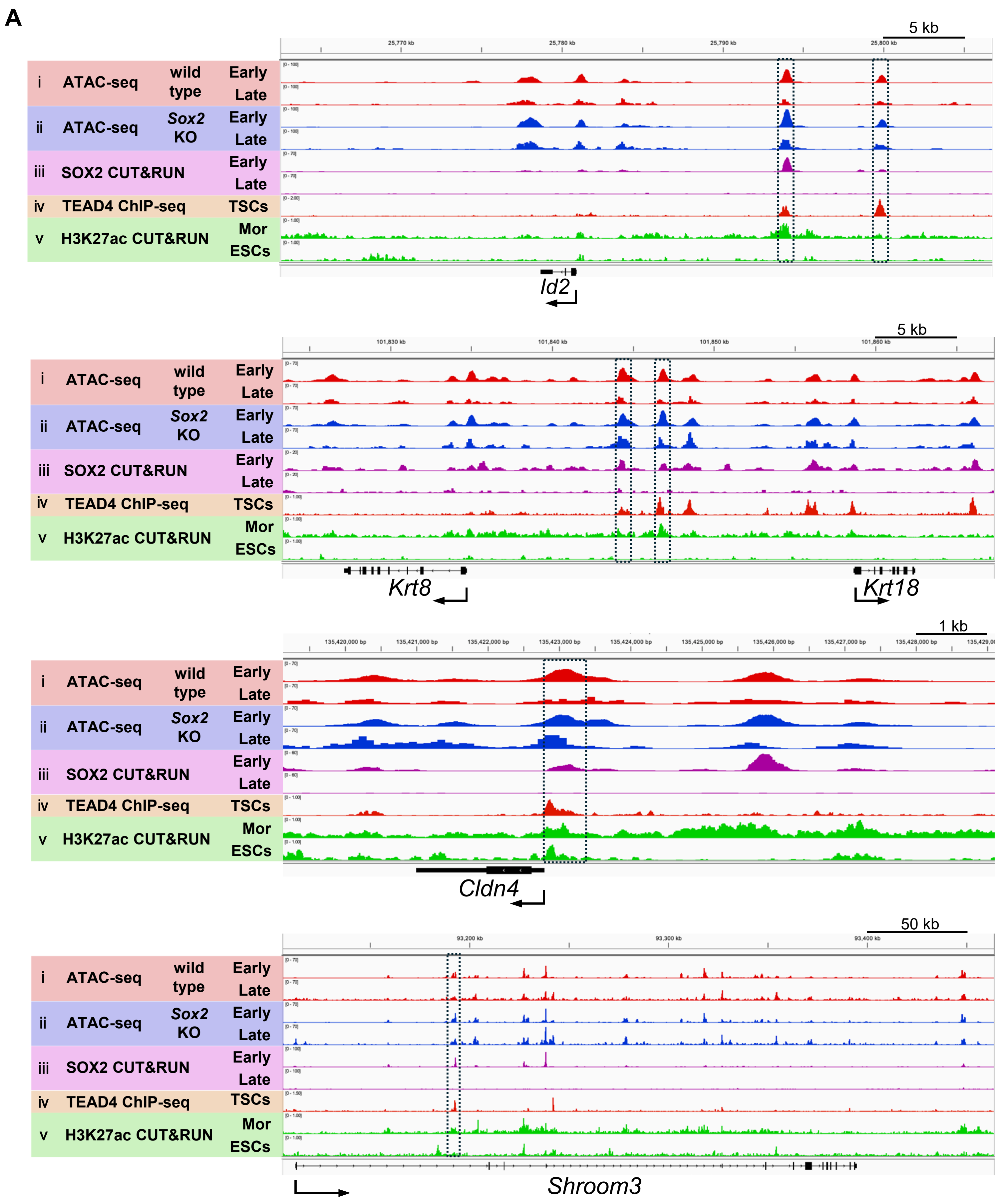

**Figure S5. SOX2 restricts the responsiveness of multiple TE genes to TEAD–YAP by remodeling chromatin accessibility, related to Figure 6.**

(A) IGV snapshots of the genomic regions surrounding TE-specific genes (*Id2* and *Krt8/18*) or apical domain genes (*Cldn4* and *Shroom3*), showing (i) ATAC-seq signals in ICMs of wild-type early and late blastocysts, (ii) ATAC-seq signals in ICMs of *Sox2 KO* early and late blastocysts, (iii) SOX2 CUT&RUN signals in early and late blastocysts, (iv) TEAD4 ChIP-seq signals in TSCs, and (v) H3K27ac CUT&RUN signals in morulae and ESCs. Dashed boxes indicate Cluster 1 high ATAC-seq peaks that overlap with SOX2 CUT&RUN peaks and TEAD4 ChIP-seq peaks in TSCs, similar to the *Gata3* TEE.

### Figure S6, related to Figure 7

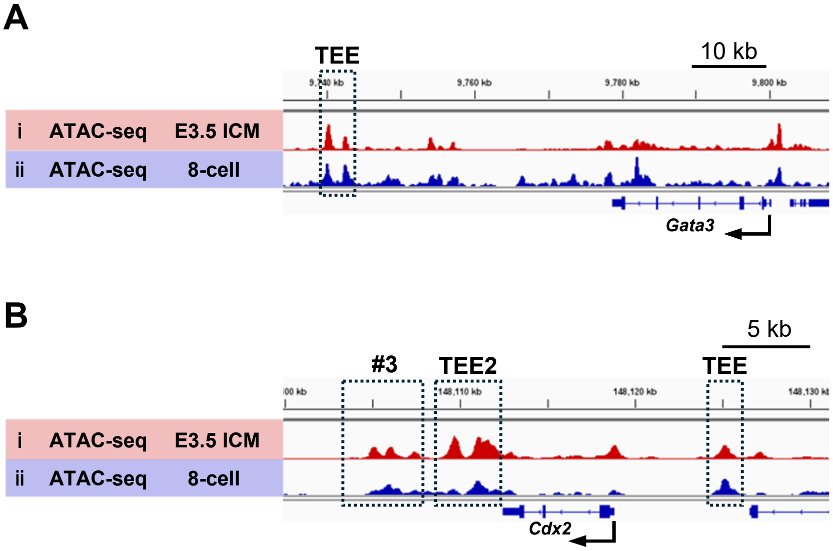


**Figure S6. TE enhancers are already accessible at the 8-cell stage, related to Figure 7**

(A) IGV snapshots of the genomic region surrounding *Gata3*, showing (i) ATAC-seq signals in ICMs of early (E3.5) blastocysts and (ii) ATAC-seq signals in 8-cell-stage embryos. The dashed box indicates TEE.

(B) IGV snapshots of the genomic region surrounding *Cdx2*, showing (i) ATAC-seq signals in ICMs of early (E3.5) blastocysts and (ii) ATAC-seq signals in 8-cell-stage embryos. The dashed boxes indicate TE enhancers: TEE[26], TEE2 and #3[54].
